# Niche specialization and cross-feeding interactions shaping gut microbial fiber degradation in a model omnivore

**DOI:** 10.64898/2026.01.22.701066

**Authors:** Rachel L. Dockman, Elizabeth A. Ottesen

**Affiliations:** Department of Microbiology, University of Georgia, Athens, Georgia, USA

**Keywords:** gut microbiome, cockroach (*Periplaneta americana*), fiber, diet, xylan, whole food, processed food

## Abstract

The gut microbiome plays an active role in host health, producing gut metabolites that influence host digestive and immune function while also mediating microbial crosstalk. Dietary fiber is a major source of important fermentation byproducts that are generally implicated in gut community stability and host wellbeing, but dissecting microbe-specific contributions to polysaccharide metabolism in the context of a complex gut community is challenging with conventional model organisms. Using the American cockroach (*Periplaneta americana*) as a model omnivore, we use chemically-defined synthetic diets to identify how complex gut microbial communities respond to two of the most abundant plant polysaccharides, xylan and cellulose. To do so, we fed cockroaches synthetic diets containing one of these fibers or a mix of both in differing ratios. Through both 16S rRNA gene profiling and RNA-seq, we show that mixed fibers enrich for organisms characteristic of the source fibers as well as additional organisms only enriched in mixed-fiber diets. Through an organism-centric pangenome approach, we identify the impact of these fibers on gut microbiome activity. We found that gut communities responded strongly to xylan, with *Bacteroidota* belonging to *Bacteroides, Dysgonomonas*, and *Parabacteroides* producing xylan-active CAZymes at high levels. Multiple groups of *Bacillota* also responded strongly to a xylan diet, but appeared to act as cross-feeding secondary degraders, producing primarily xylosidases and transcripts associated with xylose utilization. In contrast, cellulose diets were associated with higher transcriptional activity among *Fibrobacterota,* which are typically a minor component of the cockroach gut microbiome but were the primary producers of CAZymes associated with cellulose and cellobiose degradation. These experiments provide new insight into gut microbial metabolism of these complex plant polysaccharides. Further, they highlight the utility of the cockroach model and synthetic diets to answer fundamental questions about gut microbial responses to different polysaccharides alone and in combination.

## 2 Introduction

The gut microbiome exerts substantial influence over host health, conferring benefits or disadvantages depending on its composition of microbial residents and their metabolic activities within the host [1]. Therefore, understanding the factors enabling one to obtain and maintain a beneficial microbiome configuration is a key goal of microbiome research. Previous research has identified host characteristics that are central to structuring the gut microbiome, such as anatomy, habitat, and diet, as well as competitive or cooperative interactions between gut microbiota that further alter the microbiome composition [2–4]. Of these factors, diet is an especially amenable driver of gut microbiome structure, with microbial community metabolism and the resultant metabolites constrained by nutrients provided by the host. [5, 6]. There is considerable interest in leveraging individual dietary components, particularly fibers, to selectively manipulate gut microbiota into producing an optimal pool of metabolic byproducts that are conducive to human wellbeing [7–9]. However, major challenges remain in predicting and isolating microbial responses triggered by specific dietary components that occur within a complex web of microbial cross-talk and activity.

Of the macronutrients consumed by an omnivorous host, fiber is the most relevant to the gut microbiome and therefore has the most potential for selectively modulating a gut community. The term “fiber” is a broad classification encompassing any plant-derived carbohydrate that is recalcitrant to human enzymatic digestion, such as cellulose, hemicelluloses (mannan, beta-glucan, xylan), pectin, and lignin [10, 11]. While the host has first access to dietary components and can therefore extract desirable nutrients before food reaches gut microbiota, most host omnivores do not encode the cellulases or glycoside hydrolases required to break down fibers. Instead, these fibers pass directly to the gut, serving as an important source of carbon for the microbial community [12, 13]. Diets deficient in fiber disrupt microbial distribution throughout the gut [14, 15], lower diversity [16], and stimulate increased catabolism of host glycans, damaging mucus barriers protecting the gut [17, 18].

To fully understand the influence of fiber on the gut microbiome, it is vital to observe microbial responses to different fiber types. However, dietary manipulation in mammalian models is limited by the complex nutritional needs of the host. Omnivorous cockroaches such as the American cockroach (*Periplaneta americana*) are quickly emerging as model organisms for host-microbiome interactions that reflect the complexity of mammalian microbial communities while overcoming their dietary limitations [19–25].

The gut environment of cockroaches is analogous to mammals in that it can be divided into three functionally distinct sections: the foregut (crop and proventriculus) for initial amylase activity and mechanical digestion; the midgut (gastric caeca and ventriculus) for primary host digestive activity; the hindgut (colon) for waste consolidation and microbial fermentation [26–30]. Within the hindgut, cockroach gut microbiota are compositionally similar to the communities found in humans and mice, consisting primarily of the bacterial phyla *Bacteroidota*, *Bacillota*, and *Desulfobacterota* [20, 22, 24, 31]. While these bacteria are commonly found in gut communities, the niche they fill is influenced by what their host consumes. *Bacteroidota* isolates are frequently identified as primary fermenters that are capable of digesting complex polysaccharides, notably host-indigestible fibers, into their constituent molecules for either individual use or to release into the environment for consumption by secondary fermenters, such as *Bacillota* [13, 23, 32–35]. In addition to their roles in oligosaccharide scavenging, members of *Bacillota* are also associated with amino acid fermentation. Fermentation by these two phyla converts indigestible polysaccharides into metabolic byproducts such as short chain fatty acids, lactate, formate, and hydrogen, which can be absorbed by the host as vital energy sources or anaerobically respired by gut chemolithotrophs such as *Desulfobacterota* and methanogenic *Archaea* [36–38].

Previously, we have found that single fiber synthetic diets induce alterations in the gut microbiome taxonomic composition of the American cockroach despite its resilience to drastic changes in whole-food diet compositions [20, 21]. In particular, microcrystalline cellulose and xylan generated the largest differences in community structure, with the xylan-fed gut community displaying reduced diversity and fragmented co-correlation networks [21]. Cellulose and xylan are the two most abundant biopolymers found on Earth, and are therefore highly attractive subjects for fiber-based research. Cellulose is comprised of repeating glucose units bound by β(1-4) glycosidic bonds, forming a linear structure that serves as the scaffolding and backbone of plant cell walls. In contrast, xylan contains a backbone of xylose units bound by β(1-4) glycosidic bonds that can be decorated with branches of arabinose, fucose, methyl groups, acetylation, glucuronic acid, or galacturonic acid, among other sugars, serving with pectin as fortification of the cellulose scaffold [39, 40]. Both polysaccharides are indigestible to humans and cockroaches, restricting the portion of digestible carbohydrates that can be replaced in mammalian diets. Insect models do not have the same restrictions, making *P. americana* an ideal platform in which to study these fibers more deeply.

Here, we fed American cockroaches synthetic diets featuring microcrystalline cellulose, xylan, or a mixture of both as the carbon source to identify how these chemically distinct fibers influence the functional landscape of a diverse gut community. Through 16S rRNA gene sequencing paired with metatranscriptomic profiling, we observed that dominant community members exhibit large differences in their abundance and activity when confronted with these two fibers, individually and in varying mixtures. These results highlight the flexible roles played by members of a highly complex gut community living within an unfussy omnivore and provide a framework for evaluating microbial responses to distinct dietary fibers.

## 2 Materials and Methods

### 2.1 Insects

Our *Periplaneta americana* colony has been maintained in captivity at the University of Georgia for over a decade. Mixed age and sex stock insects are maintained at room temperature in glass aquarium tanks with wood chip bedding and cardboard tubes for shelter in a 12:12 light:dark cycle. Water via cellulose sponge fit to a Tupperware reservoir and dog chow (Purina ONE chicken & rice formula, approximately 26% protein, 16% fat, and 3% fiber) are provided to stock colonies *ad libitum*.

### 2.2 Diet Composition

Synthetic diets were prepared as described in [21] and briefly summarized here. The synthetic diets contained, as percent dry weight, 0.5% Vanderzant vitamin mix (catalog #: 903244, MP Biomedicals, USA), 3% Wesson salt mix (catalog #: 902851, MP Biomedicals), 8% peptone (catalog #: J636, Amresco, VWR International, USA), 17% casein (catalog #: C3400, Sigma-Aldrich,USA), and 1% cholesterol (catalog #: 0433, VWR). The remaining 70.5% consisted of either xylan from corn core (catalog #TCX0078, TCI Chemicals, USA), microcrystalline cellulose (MCC; 51um particle size; catalog #435236, Sigma-Aldrich), or a mixture of these carbohydrates in ratios of 1:1, 3:1, or 6:1. Throughout, ratio diets will referred to by the percentage of the majority-fiber. Diets with 6:1 ratios will be referred to as 86% “cellulose” or “86% xylan”, 3:1 will be “75% cellulose” or “75%” xylan, and 1:1 will be referred to as “50% mix”. For each diet, dry components were suspended in sufficient volumes of diH_2_O to form a homogenous dough, shaped into pellets, then dehydrated at 65°C until solid. Food pellets were stored at −20°C until use.

### 2.3 Experimental Design

As described in [21] and [20], healthy mixed-sex *P. americana* adults were selected from the stock colony and moved to experimental tanks containing bleach-sterilized pebbles and polyvinyl chloride tubes for footing and shelter, with synthetic diets and water provided *ad libitum* in glass petri dishes. Dietary experiments lasted for two weeks, during which food and water were replaced daily and visible debris or oothecae were removed as needed.

All insects were sacrificed for sample collection upon completion of the two-week treatment period via cold-immobilization, decapitation, and dissection. The insects were dissected on individual clean culture plates with fine tipped forceps to remove exoskeleton and fat body adhered to the gut wall. The cleaned gut was then transferred to an aluminum dish on dry ice for freezing and divided into foregut, midgut, and hindgut sections. Hindgut sections to be used for DNA extraction were collected into 500uL phosphate-buffered saline (1X PBS), disrupted with a sterile pestle to suspend gut-attached bacteria, and stored at –20°C until DNA extraction.

To construct host transcriptomes and hindgut metatranscriptomes, five samples were collected from each individual insect: hindgut lumen, midgut lumen, hindgut tissue, midgut tissue, and fat body. The insects were dissected as described above, with the addition of gut-adhered visceral fat body collection while cleaning the gut surface. All samples for RNA sequencing were stored in 100uL RNAlater (Invitrogen, Thermo Fisher, USA) immediately upon removal from the dry ice. Fat body samples were directly deposited during the dissection. Midgut and hindgut sections were transferred to a sterile 1.5mL tube and pressed against the tube wall with a pestle to coax out gut lumen contents without pulverizing the tissues, after which the sample was vortexed in 100uL RNAlater. The pestle was again used to lightly wring out remaining lumen contents from the gut tissue, then to transfer the empty guts into fresh 1.5mL tubes containing RNAlater. Samples destined for RNA-seq were stored at −80°C until use.

### 2.4 Extraction and Sequencing of 16S rRNA Gene Libraries

DNA was extracted from 200µL aliquots of individual samples using the EZNA Bacterial DNA Kit (Omega Biotek, Norcross, GA, USA) as described in detail in [21]. The extracted DNA, suspended in 50uL of EZNA elution buffer, was checked for quality and quantitated using the Take3 plate for BioTek plate readers (Agilent, USA) prior to barcoding and Illumina-compatible library preparation.

The V4 region of the 16S rRNA gene was amplified and appended with barcoded primers from 5-10ng DNA per sample via 2-step polymerase chain reaction (PCR) using 0.02U/L Q5 Hot Start high-fidelity DNA polymerase (New England BioLabs, Ipswich, MA, USA), described in [20, 24, 41]. The final products were checked via gel electrophoresis and cleaned as outlined in the Omega Biotek Cycle Pure kit, then pooled at equimolar proportions for sequencing. The prepared library was sequenced with 250 base pair paired-end Illumina MiSeq sequencing at the Georgia Genomics and Bioinformatics Core at the University of Georgia.

### 2.5 RNA Extraction

Both host and microbial RNA were extracted using the EZNA Total RNA Kit II (Omega-Biotek), which combines the phenol/guanidine isothiocyanate-based RNA-Solv Reagent (Omega-Biotek) with column-based nucleic acid purification. While both sample types shared most steps in RNA extraction, they differ in homogenization method. For host transcriptomes, half of each gut wall sample (split lengthwise) or approximately 15mg fat body was homogenized directly in 1mL RNA-Solv with an Ultra-Turrax rotor-stator homogenizer (Janke & Kunkel, Germany). The homogenized samples were phase-separated with 200uL chloroform (VWR) and the aqueous phase column-purified as described in the kit instructions. The final product was eluted into 50uL nuclease-free water.

For microbial RNA extraction, 50uL of each sample was vortexed with 200uL ice-cold RNase-free PBS, then centrifuged at 5,000g for five minutes to pellet. The supernatant was removed, and the pellet resuspended in 50uL of a 30mg/mL lysozyme/RNase-free TE buffer solution with 4uL Superase-IN RNase inhibitor (Invitrogen) via 30s of vortexing. Samples were incubated in a shaking incubator (Eppendorf, USA) for 10 minutes at 30°C and 350rpm, then transferred to a screw-top 2mL tube containing 50mg 0.1mm silica beads (BioSpec, USA) and 1 mL RNA-Solv. Cell lysis was performed via four cycles of bead beating at max speed on a vortex for 30s followed by 30s on ice, after which samples were phase-separated with 200uL chloroform and RNA isolated as above.

Isolated RNA from all sample types were incubated for 20 min at 37°C with 1uL TURBO DNase enzyme (Invitrogen) and 5uL TURBO DNase reaction buffer to destroy any contaminating DNA, then purified using the column-based EZNA MicroElute RNA Clean-Up Kit (Omega-Biotek). Microbial RNA was eluted once in 15uL nuclease-free water, while host samples were eluted twice for 30uL total of purified RNA per sample. The RNA content was quantitated via the Take3 plate, then assessed for quality with the Agilent Bioanalyzer, and samples were stored at −80°C until library preparation.

### 2.6 RNA Library Construction

Host tissue mRNA was isolated with the NEBNext® Poly(A) mRNA Magnetic Isolation Module (NEB) according to kit instructions, while prokaryotic gut lumen RNA was ribodepleted with biotinylated probes constructed from the extracted DNA of pooled MCC-fed and xylan-fed cockroach guts based on the methods described in [22, 42]. Briefly, T7-promoter appended primers were used for PCR to amplify genes corresponding to both bacterial and archaeal 16S and 23S rRNA, in addition to cockroach 18S, 28S, and ITS regions (**Supplemental Table 1**). The purified PCR products were then transcribed with the AmpliScribe T7-Flash Biotin-RNA Transcription kit (Lucigen Corp, Middleton, WI) and cleaned with the MEGAclear Transcription Clean-up Kit (Invitrogen). These probes were hybridized with total prokaryotic RNA in saline-sodium citrate (SSC) buffer and formamide, then captured with streptavidin magnetic beads (NEB) which had been cleaned with 0.1N NaOH and SSC buffer ahead of time. The mRNA remaining in the supernatant was cleaned with the EZNA RNA clean-up kit prior to library preparation.

Host and microbial mRNA libraries were constructed using the NEBNext® Ultra II Directional RNA Library Prep Kit for Illumina (NEB), following instructions to obtain 300 bp inserts flanked by Illumina-compatible index primers (NEBNext® Multiplex Oligos for Illumina®). Following size and quality verification with the Bioanalyzer, the barcoded samples were pooled together and sent to Novogene (Sacramento, CA, USA) for Illumina NovaSeq sequencing.

### 2.7 16S rRNA Bioinformatics

The amplicon data collected in this study was processed in R (version 4.2.1) using the R package DADA2 (version 1.24.0), with Amplicon Sequence Variants (ASVs) previously obtained from cockroach gut 16S rRNA gene sequencing included as a priors table [21, 43, 44]. Taxonomy was assigned using DADA2 and the ARB Silva v138 classifier to the species level, followed by filtering to remove sequences matching eukaryotic (chloroplast, mitochondria) or endosymbiotic *Blattabacterium* DNA [43, 45].

Alpha and beta diversity analyses were performed on rarefied count tables (10,790 reads) with the R package vegan (version 2.6-4) [46]. Alpha diversity was calculated for Shannon index (*diversity()*), number of observed ASVs, and Pielou’s evenness (calculated as Shannon/log(Observed)) and evaluated for significance with Wilcoxon rank sum test (pairwise comparisons; *pairwise.wilcox.test()*) and Kruskal-Wallis test (multi-group comparisons; *kruskal.test()*). Bray-Curtis dissimilarity and Sørensen index matrices were determined with *vegdist()* then ordinated using non-metric multidimensional scaling (NMDS; function *metaMDS*). Overall ordination quality was assessed with PERMANOVA (function *adonis2(*)), and *envfit()* was used to fit “dietary xylan percent” to the plots.

Differential abundance analysis of the ASVs was conducted using DESeq2 (version 1.36) with parameters ‘fitType = “local”’ and ‘design = ∼ Diet’ [47]. Pairwise result tables were extracted for all diet comparisons and filtered to retain enriched ASVs with an adjusted p-value < 0.05 and baseMean > 100. Overlap in ASVs enriched by single- and mixed-fiber diets was determined by designating ASVs enriched in any of the four ratio diets as enriched by mixed-fiber, then comparing these ASVs to those enriched on the two single-fiber diets.

### 2.8 Metatranscriptome Processing and Pangenome Construction

The paired-end metatranscriptome sequencing data were processed on the UGA computing cluster using both established programs and custom in-house perl scripts. Joint Genome Institute (JGI) programs BBduk and BBSplit (https://jgi.doe.gov/data-and-tools/software-tools/bbtools/) were used for adapter removal and contamination filtering respectively; BBSplit indexes contained the genome and plasmid of the blattid endosymbiont, *Blattabacterium* [48], and host sequences were flagged and removed based on alignment to the *P. americana* reference genome available on NCBI (BioProject PRJNA1098420) [49]. The remaining reads were screened for rRNA contamination with SortMeRNA and paired mRNA reads were merged using the JGI program BBMerge [50, 51]. Any paired reads that could not be merged using BBMerge were combined with a 10-N spacer sequence using an in-lab script. The merged consensus reads were repaired and filtered with a BBMap shell script (repair.sh), then translated with prodigal [52] for compatibility with DIAMOND blastp [53].

Alignment of the translated reads was performed against the nonredundant bacterial and archaea RefSeq protein databases [54, 55], in addition to a custom database comprising single cell genomes derived from the *P. americana* hindgut [22]. Results from these two DIAMOND runs were combined with an in-lab Perl script to maintain higher scoring hits and improve specificity of our annotations to our host organism. **Supplemental Table 2** reports per-sample reads remaining following each processing step.

To minimize the impact of the limited and partial reference genomes available for many cockroach gut microbiota, we mapped transcripts to approximately genus-level pangenomes for key cockroach gut taxa as described in [56]. Pangenomes were constructed with anvi’o, which aligns and classifies input proteins from user-selected genomes to determine gene clusters that share structural and functional features [57–61]. Gene clusters were annotated with clusters of orthologous genes (COGs) [62, 63], KEGG orthologs [64–66], and carbohydrate-active enzymes (CAZymes) [67–69] during the anvi’o workflow. The final summary file generated in the anvi’o pipeline contains the computed gene clusters that are annotated with protein accessions from the reference genomes, COGs, KOs, and CAZymes. In-lab Perl scripts were used to match hits in the DIAMOND blastp output to the gene clusters and build pangenome count tables for further analysis.

Twelve previously constructed pangenomes (*Bacteroides, Parabacteroides, Dysgonomonas, Alistipes, Odoribacter, Paludibacteraceae, Clostridiaceae, Oscillospiraceae group 1, Oscillospiraceae group 2, Enterococcaceae, Desulfovibrio,* and *Desulfosarcina)* that were designed based on cockroach gut members [70] were used in this study, as well as five new pangenomes: *Lachnospiraceae A, Lachnospiraceae B, Lachnospiraceae C, Fusobacterium*, and *Fibrobacterota.* Since RNAseq can poorly differentiate *Bacillota* members, co-occurrence patterns observed in this experiment were considered when constructing these pangenomes. *Lachnospiraceae* groups were determined based on Kendall rank correlations calculated for the taxonomic abundance of *Lachnospiraceae* genera with absolute abundances > 1% across ratio diets (**Supplemental Figure S1**). Genomes that were used as references to create these pangenomes are listed in **Supplemental Table 3.**

### 2.9 Metatranscriptome analyses

Before analyzing transcripts in the context of pangenomes, the overall taxonomic and structural composition of the metatranscriptomes was assessed based on the top-scoring DIAMOND hits (based on highest bit scores) per read. Since many reads had multiple top-scoring hits, the count of taxonomic identities per read was split to allow each possibility to be considered, thus creating a weighted count table that acknowledged all possible microbes present. These weighted counts were rarefied to the smallest sample size (4,738,232) and used as input into the vegan R package for diversity calculations (alpha: *diversity();* beta: *vegdist()*), ordination (*metaMDS()*), and statistical analyses (*adonis2(), kruskal.test(), pairwise.wilcox.test()*) as described for the 16S rRNA amplicon analysis.

Pangenomes were analyzed for diet-related community structure using gene cluster count tables. Enrichment was determined using DESeq2, after which the number of significant gene clusters were extracted for each pairwise comparison and visualized as a heatmap. Principal component (PCA) and redundancy analyses (RDA) were performed on count tables after transformation with the DESeq2 variance stabilizing transformation function *vst()*. The function *rda()* from the vegan package was used for both; when no model is selected, the function generates unconstrained PCs. Constrained models used for RDA include percent xylan (numeric), whether the diet contains xylan in any amount (“Contains xylan”), whether the diet contains cellulose in any amount (“Contains cellulose”), and “Diet”. Component values were obtained with vegan::*summary()* and model significance was assessed with ANOVA (function *anova()*).

The carbohydrate degrading capacity of the pangenomes was analyzed based on gene clusters with CAZyme assignments. Gene clusters sharing the same CAZyme assignment were aggregated together; if a gene cluster had more than one associated CAZyme, it was overcounted to consider both assignments. Substrate specificity for the various CAZyme families was annotated using a mapping file “dbCAN-sub.substrate.mapping.xls” available from the dbCAN2 server [67], with a subset replicated in **Supplemental Table 4**. Patterns of CAZyme expression associated with xylan or cellulose was visualized as a heatmap of overall read abundance rather than scaling by samples or CAZyme family.

Metabolic enrichment was determined via DESeq2 for the pangenomes using the individual KEGG orthologs assigned to gene clusters as well as gene clusters collapsed into their level 3 assignments for carbohydrate and amino acid metabolism pathways [64–66]. Enrichment in total pathway expression between the 100% xylan and cellulose diets was displayed as a color-block heatmap. For comparison across the ratio diets, pathways were visualized with heat map bubble plots sized by relative abundance of the entire sample transcript count and color scaled by relative abundance within the pangenome. Pangenome enrichment of individual enzymatic steps were annotated onto KEGG metabolic maps using the R package pathview [71].

## 3 Results

### 3.1 Mixed fiber diets produce intermediary 16S rRNA profiles between pure xylan and cellulose diets

To determine how cellulose and xylan influence the composition of the cockroach gut microbiome, we prepared six synthetic diets featuring xylan, microcrystalline cellulose, or a mix of these two fibers with a fixed basee of peptone, casein, micronutrients, and cholesterol as described in previous work [21]. Mixed fiber diets contained xylan and cellulose in ratios of 6:1, 3:1, 1:3, and 1:6 by weight. These six diets were fed to mixed-sex adult cockroaches (n=10/diet) for two weeks, after which we sequenced the 16S rRNA gene profile of their hindgut microbiomes. In total, 2,715,590 quality- and contaminant-filtered reads were obtained from 60 samples with an average read depth of 45,259 (range: 7,656 – 130,911). Taxonomic assignment via the SILVA (v138.1) 16S rRNA database classified most reads as *Bacillota*, *Bacteroidota*, or *Desulfobacterota* (**Supplemental Figure S2**), consistent with previous observations of cockroach gut taxonomic composition [20, 21, 25].

We examined the impact of dietary fiber composition on the alpha and beta diversity of the cockroach hindgut microbiome. Overall, host diet influenced sample alpha diversity in terms of Shannon index (Kruskal-Wallis, p < 0.001), richness (Kruskal-Wallis, p <0.01) and evenness (Kruskal-Wallis, p < 0.001) **(Figure 1A-C)**. Interestingly, pairwise analyses between the mixed-fiber and 100% cellulose diets found no direct differences. Only diets containing 100% xylan produced gut communities with notably lower alpha diversity than mixed-fiber or 100% cellulose diets (Wilcoxon paired tests, p < 0.05 each; **Figure 1A-C),** suggesting that replacing even 25% of xylan with cellulose in these synthetic diets increased gut microbial diversity and stability. Similarly, we found that dietary fiber was strongly correlated with microbiome beta diversity in both abundance-weighted Bray-Curtis (PERMANOVA: R2 = 0.271, p. < 0.001; **Figure 1D)** and abundance-independent Sørensen index (PERMANOVA: R2 = 0.205, p. < 0.001; **Figure 1E**) NMDS ordinations. We noticed that, rather than forming discrete diet-dependent clusters, mixed-fiber samples appeared to disperse from 100% cellulose samples towards 100% xylan samples in an overlapping, gradient-like manner. Indeed, fitting xylan percentage to the ordinations with envfit confirmed an additive effect of increasing xylan proportion to community structure similarity on both abundance-weighted (r=0.627, p < 0.001; **Figure 1D**) and unweighted (r=0.358, p < 0.001; **Figure 1E**) measures.

**Figure 1:**
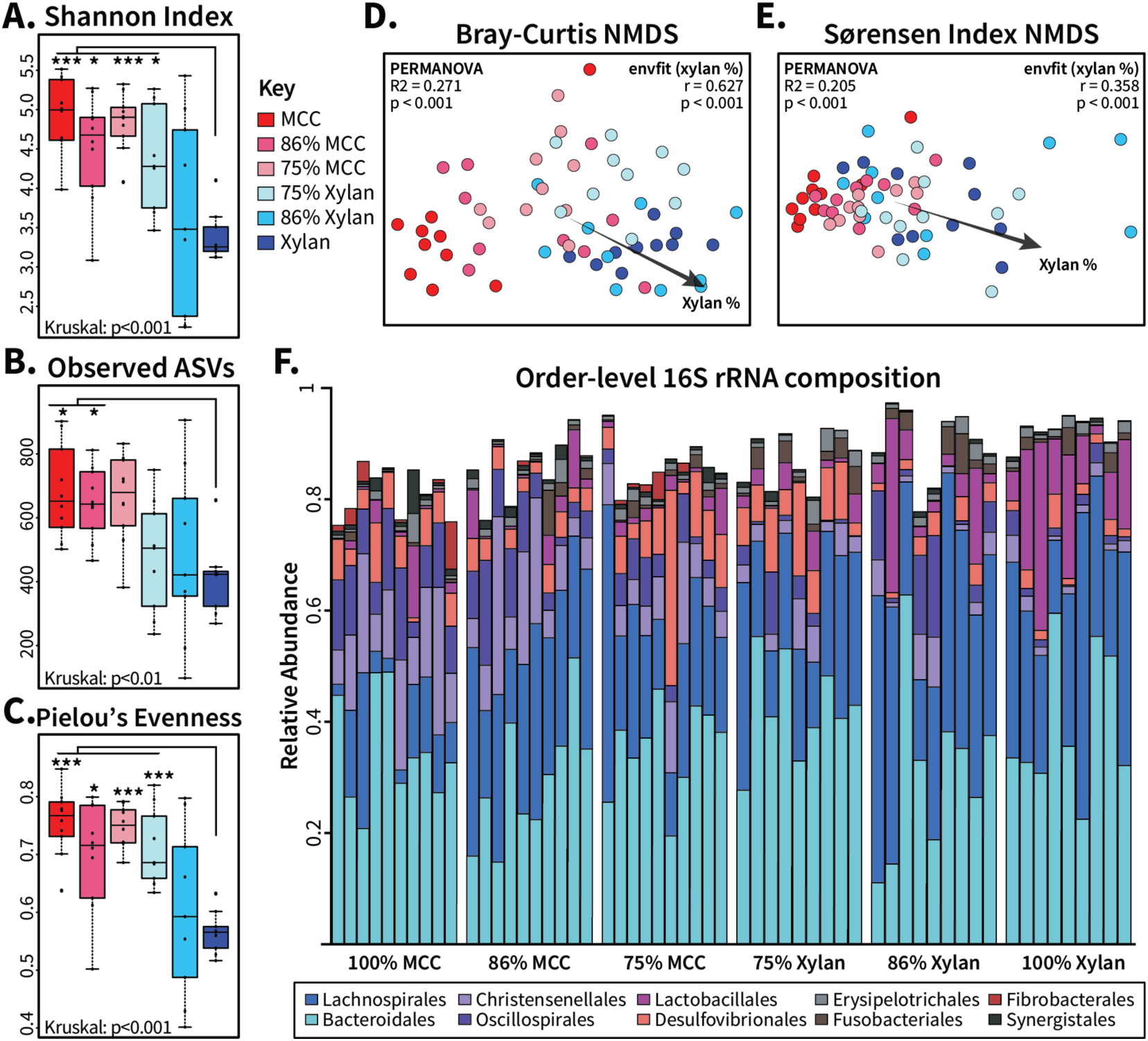
Community composition of gut microbiota from ratio diets show dose-dependent shift from cellulose-fed communities to xylan-fed communities. Samples were rarefied to 10,790 reads prior to calculation of (A) Shannon index, (B) richness via observed ASVs, and (C) Pielou’s evenness and visualized by box plots. Significance was determined by Kruskal-Wallis test and Wilcoxon test for group and pairwise comparisons, respectively. Ordinations of the rarefied samples were obtained with beta diversity measures (D) Bray-Curtis dissimilarity and (E) Sørensen index. PERMANOVA was applied to determine the overall significance of diet in ordination structure with “percent xylan” fit to the data and analyzed using envfit. (F) ASV count tables were aggregated at the order taxonomic level to calculate relative abundance, and the top ten most abundant bacterial orders were visualized. MCC: microcrystalline cellulose; * = p < 0.05; ** = p < 0.01; *** = p < 0.001

These fiber-dependent differences in microbiome diversity were reflected in the taxonomic composition of the cockroach gut microbiota. As in our previous observations [21], 100% xylan and 100% cellulose diets selected for divergent gut microbial taxonomic compositions at the order level; mixed fiber diets seemed to form intermediate stages between the pure diets (**Figure 1F**). Taxonomic signatures such as *Christensenelalles* and *Oscillospirales* in the cellulose diet, for example, were diminished but still evident in the mixed-fiber diets. Likewise, *Lachnospirales* and *Lactobacillales* abundant in the 100% xylan diet also formed a larger proportion of the mixed communities than in the cellulose diet. To identify potential organisms driving these large-scale taxonomic shifts, we applied DESeq2 to the amplicon sequence variant (ASV) count table with a baseMean cut-off of 100 and adjusted p < 0.05. In total, 84 ASVs differed in at least one pairwise comparison between the various diets **(Supplemental File A)**. Comparing xylan to cellulose produced the most (68) differentially abundant ASVs of all pairwise comparisons, while the 100% cellulose and xylan diets shared more enriched ASVs with their corresponding fiber-majority diets than their fiber-minority diets (**Supplemental Figure S3A**). When these diet-responsive ASVs were placed in a Venn diagram based on enrichment by xylan, cellulose, any mixed fiber ratio, or multiple diet conditions (**Supplemental Figure S3B)**, most ASVs appeared in the intersections of one pure fiber and the mixed fibers. Interestingly, we found that the 100% fiber diets uniquely enriched for one organism each (cellulose: *ChristensenellaceaeR7.5;* xylan: *Bacteroidales.0*) in contrast to the mixed-fiber diets, which enriched for 15 ASVs including five *Lachnospiraceae* and four *Alistipes* members. (**Supplemental Figure S3B**). Charting the log2 fold changes between all ASVs enriched by xylan (**Supplemental Figure S3C**) or cellulose (**Supplemental Figure S3D**) revealed that many enriched ASVs displayed similar gradient-like abundance differences as sample-wide diversity measures found, especially in comparison to the cellulose diet. However, many ASVs enriched by xylan and cellulose belonged to the same lineage. Within *Bacteroides*, for example, ASVs were enriched both by xylan (7) and by cellulose (4), suggesting species-level differences in microbiome composition missed by 16S amplicon analyses.

### 3.2 Taxonomic composition of metatranscriptomes

In order to establish the functional basis through which fiber source modulates the cockroach gut microbiome, we gave adult insects one of five synthetic diets containing xylan, cellulose, or a mix of both fibers (ratios of 6:1 1:1, 1:6) and obtained 30 (n=6/diet) individual paired-end hindgut microbiota metatranscriptome libraries. In total, 4,552,148,064 hindgut lumen reads were recovered (**Supplemental Table 2**). After filtering for contamination and read quality, a total of 614,662,256 consensus pairs (range: 6,013,230- 37,692,726 per sample) were obtained. Translation and alignment of these pairs to non-redundant bacterial and archaeal RefSeq protein databases and cockroach-derived single cell genomes produced 484,022,581 successfully annotated proteins (78.7% of paired reads, 4,740,617-31,943,483 per sample).

In general, the taxonomic composition obtained from the RefSeq annotations of aligned metatranscriptomic reads reflected phylum-level distribution patterns found in 16S amplicon data generated from the same fiber sources (**Supplemental Figure S2**). Compared to xylan, the 100% cellulose diet produced metatranscriptomes with higher alpha diversity **(Supplemental Figure S4A-C)** and formed denser clusters during NMDS ordinations of Bray-Curtis dissimilarity (**Supplemental Figure S4D**). At finer taxonomic resolutions, differences arose between microbial composition of the amplicon and metatranscriptomic datasets. For example, at the order-level (**Figure 2A**), *Bacteroidales* activity was strongly associated with increasing xylan percentage, with an average relative abundance of 63% compared to 35% observed in the cellulose diet; 16S amplicon sequencing captured a much smaller difference of 39% in xylan vs 34% in cellulose. *Lactobacillales,* which comprised an average 14% of xylan 16S amplicon reads, showed little activity in the metatranscriptomes, staying under 1% of transcripts for the single fiber diets. These discrepancies suggest that some community members have activity levels disproportionate to their overall abundance, further complicating the dynamics underlying microbiome-fiber interactions.

**Figure 2:**
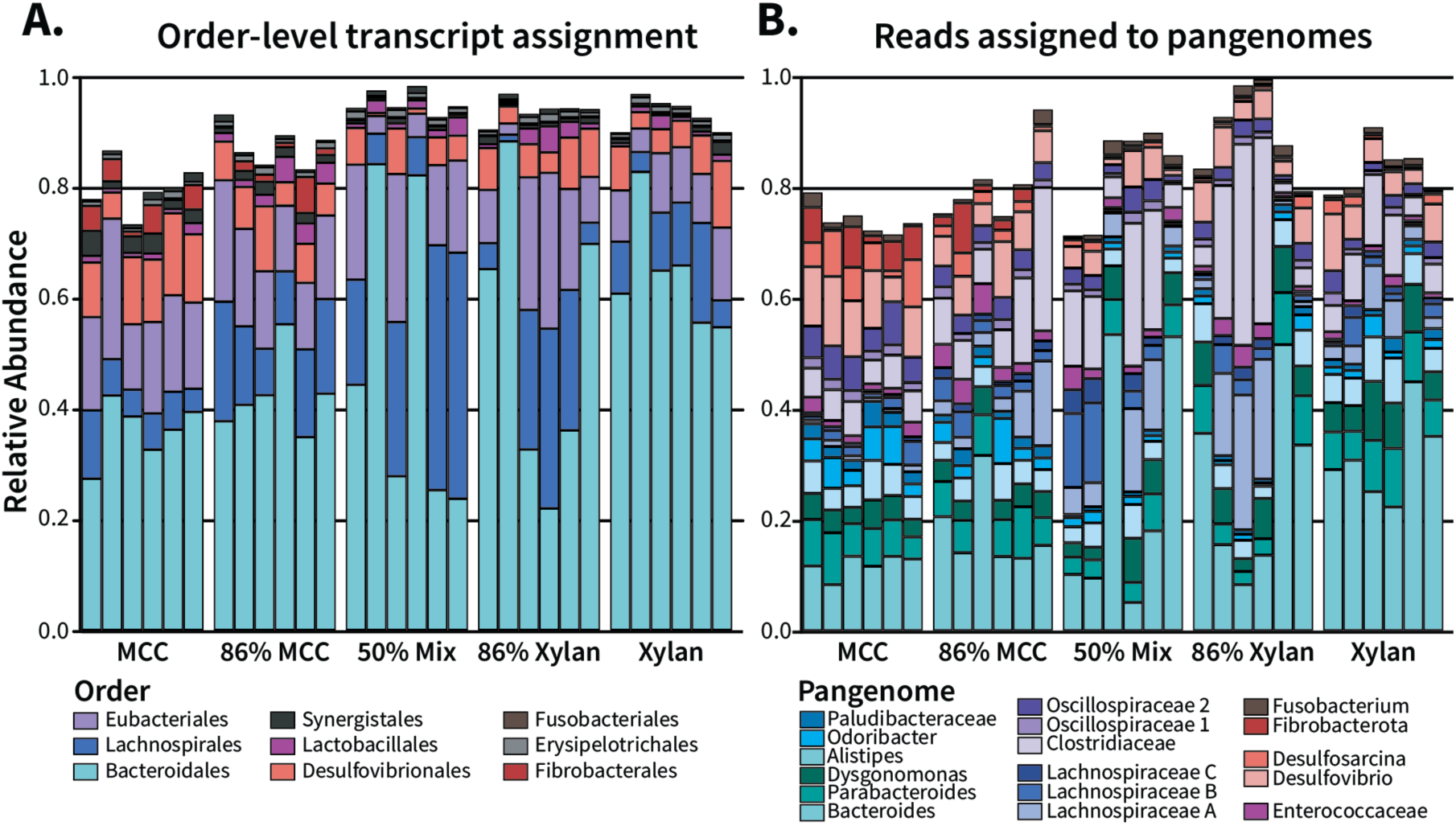
Pangenome assignment captures the most abundant taxonomic groups identified in RNA annotation. **(A)** Count tables tabulating the taxonomic composition of weighted RefSeq and single-cell genome hits were collapsed at the order level for relative abundance calculation. Relevant orders with high abundance in the previous 16S rRNA gene experiments were selected for visualization. (**B**) Relative abundance of metatranscriptomic reads matched to the 17 pangenomes.

### 3.3 Pangenome Overall Comparisons

To facilitate transcriptional analyses of cockroach-associated microbes, we used an in-house pipeline to map transcripts to 17 pangenomes representing key taxa within the cockroach gut microbiota [22, 56, 57]. These include 12 previously constructed pangenomes (*Bacteroides, Parabacteroides, Dysgonomonas, Alistipes, Odoribacter, Paludibacteraceae, Clostridiaceae, Oscillospiraceae group 1, Oscillospiraceae group 2, Enterococcaceae, Desulfovibrio,* and *Desulfosarcina)* and five additional pangenomes (*Lachnospiraceae A, Lachnospiraceae B, Lachnospiraceae C, Fusobacterium*, and *Fibrobacterota*) that were newly constructed for this analysis. Altogether, between 66.5% and 94.7% of translated reads per sample were assigned to a pangenome (**Figure 2B).** The largest portion of reads were mapped to *Bacteroides* followed by *Clostridiaceae, Lachnospiraceae A,* and *Desulfovibrio,* while *Fibrobacterota* had the fewest (**Table 1**). On average, we annotated 78.6% of reads per pangenome with KEGG orthologs (KO) and 4.6% of reads per pangenome with carbohydrate active enzymes (CAZymes) (**Table 1**).

**Table 1:** Reads mapped to pangenome gene clusters and the percent that were assigned KEGG orthologs or CAZyme annotations.

|  | Pangenome | Reads mapped | Reads with KO (%) | Unique KO terms | Reads with CAZyme (%) | Unique CAZymes |
| --- | --- | --- | --- | --- | --- | --- |
| *Bacteroidota* | *Alistipes* | 20,992,939 | 78.47 | 1897 | 3.41 | 193 |
|  | *Bacteroides* | 110,224,278 | 74.31 | 1572 | 7.26 | 172 |
|  | *Dysgonomonas* | 25,551,436 | 75.98 | 1902 | 11.27 | 207 |
|  | *Odoribacter* | 8,493,413 | 77.83 | 1535 | 3.17 | 84 |
|  | *Paludibacteraceae* | 5,653,004 | 82.97 | 1465 | 4.91 | 144 |
|  | *Parabacteroides* | 28,204,621 | 73.52 | 1757 | 8.36 | 201 |
| *Bacillota* | *Clostridiaceae* | 65,006,931 | 81.07 | 3473 | 3.18 | 255 |
|  | *Enterococcaceae* | 8,488,768 | 68.85 | 2398 | 7.06 | 127 |
|  | *Lachnospiraceae A* | 42,841,465 | 78.57 | 2807 | 2.99 | 197 |
|  | *Lachnospiraceae B* | 26,114,106 | 84.87 | 2763 | 4.04 | 253 |
|  | *Lachnospiraceae C* | 16,067,332 | 85.95 | 2854 | 2.84 | 213 |
|  | *Oscillospiraceae 1* | 5,007,584 | 74.43 | 1820 | 6.13 | 94 |
|  | *Oscillospiraceae 2* | 13,510,807 | 81.80 | 2743 | 3.21 | 139 |
| Other | *Desulfovibrio* | 26,612,778 | 81.67 | 2935 | 2.00 | 107 |
|  | *Desulfosarcina* | 6,531,593 | 79.45 | 2438 | 1.44 | 63 |
|  | *Fibrobacterota* | 3,380,195 | 75.79 | 1249 | 4.84 | 82 |
|  | *Fusobacterium* | 4,780,018 | 79.89 | 1757 | 1.39 | 43 |

Principal components ordination of the pangenome transcriptomes showed that communities from the 100% cellulose diet formed a distinct cluster in the *Bacteroidota* genera *Bacteroides* and *Alistipes* **(Supplemental Figure S5A)** and all *Bacillota* pangenomes (**Supplemental Figure S5B).** The remaining MCC-*Bacteroidota* populations displayed some overlap with the majority cellulose diet, while *Fibrobacterota*, which stratified more variation along the first PC (64.5%) than any other pangenome, clustered the 86% and 100% cellulose diets together far away from other samples **(Supplemental Figure S5C)**. This aligns well with pairwise analyses of differential gene expression between diets, which frequently found that the largest number of differentially expressed gene clusters were identified in pairwise analyses between cellulose alone vs. diets containing any amount of xylan (**Supplemental Figure S6).**

Redundancy analysis (RDA) from the R package vegan was applied to identify how well transcriptome variance was explained by dietary factors [46]. Predictably, stratifying by “diet” as a factor explained the most variation for all pangenomes but especially for *Fibrobacterota* (53.7%) and *Bacteroides* (29.85%), although constrained ordination did not capture the entirety of variation explained by unconstrained PCA (**Supplemental Figure S5)**. We also tested models based on the presence vs absence of the fibers (e.g. 100% cellulose vs all other diets) and a numeric model of percent xylan. We observed phylum-dependent differences in how the models fit each pangenome; all *Bacillota* were more strongly separated (or equally separated for the *Oscillospiraceae* groups) by the addition of xylan in any amount to a diet, while all other pangenomes were better described by the numeric percent of xylan the diets contained (**Table 2**). Despite this difference, all pangenomes shared an interesting pattern: the presence or absence of xylan explained far more variation than that of cellulose, which only explained significant variation in *Enterococcaceae* (ANOVA: F = 2.19, p < 0.01) and *Fibrobacterota* (ANOVA: F = 4.03, p < 0.05). Overall, these results indicate there is a greater impact of xylan on the activity of gut microbiota that manifests in a phylum-specific manner.

**Table 4.2:** Redundancy analysis of pangenome gene cluster expression.

|  | Model Factor | Diet | % xylan | Contains xylan | Contains cellulose | Unconstrained PC1 |
| --- | --- | --- | --- | --- | --- | --- |
| *Bacteroidota* | Alistipes | 15.5% | 12.6% | 12.2% | 4.5%<sup>n.s.</sup> | 24.1% |
|  | Bacteroides | 29.9% | 26.3% | 21.7% | 6.9%<sup>n.s.</sup> | 38.8% |
|  | Dysgonomonas | 15.9% | 13.2% | 12.6% | 5.4%<sup>n.s.</sup> | 25.2% |
|  | Odoribacter | 18.5% | 16.4% | 11.4% | 5.5%<sup>n.s.</sup> | 30.7% |
|  | Paludibacteraceae | 16.8% | 12.5% | 10.9% | 4.7%<sup>n.s.</sup> | 24.9% |
|  | Parabacteroides | 16.8% | 13.7% | 12.5% | 4.8%<sup>n.s.</sup> | 28.2% |
| *Bacillota* | Clostridiaceae | 25.4% | 16.9% | 23.0% | 4.3%<sup>n.s.</sup> | 31.1% |
|  | Enterococcaceae | 15.9% | 10.6% | 15.2% | 7.2% | 21.2% |
|  | Lachnospiraceae A | 28.0% | 18.1% | 26.7% | 4.6%<sup>n.s.</sup> | 31.1% |
|  | Lachnospiraceae B | 17.6% | 13.0% | 16.2% | 4.4%<sup>n.s.</sup> | 21.5% |
|  | Lachnospiraceae C | 19.3% | 13.7% | 17.7% | 4.7%<sup>n.s.</sup> | 22.5% |
|  | Oscillospiraceae 1 | 17.2% | 13.8% | 13.8% | 4.6%<sup>n.s.</sup> | 21.4% |
|  | Oscillospiraceae 2 | 18.4% | 14.3% | 14.3% | 4.7%<sup>n.s.</sup> | 22.4% |
| Other | Desulfovibrio | 11.2% | 9.1% | 7.8% | 4.8%<sup>n.s.</sup> | 19.3% |
|  | Desulfosarcina | 15.0% | 11.7% | 8.2% | 3.6%<sup>n.s.</sup> | 35.9% |
|  | Fibrobacterota | 53.7% | 46.9% | 22.7% | 12.6% | 64.5% |
|  | Fusobacterium | 13.0% | 10.9% | 9.7% | 5.1%<sup>n.s.</sup> | 17.8% |

### 3.4 Changes in metabolic gene expression across diets

To examine the metabolic impacts of diets containing xylan or cellulose as their only carbohydrate source, we aggregated transcripts assigned to KEGG orthologs by KEGG metabolic pathway and analyzed the differential expression of carbohydrate (**Figure 3A**) and protein (**Figure 3B**) pathways. Only three pathways were enriched exclusively by xylan (galactose degradation, pentose interconversions) or cellulose (C5-branched dibasic acid metabolism); rather, fibers induced organism-specific differences in pathway expression (**Figure 3**). However, within *Bacillota,* we observed a consistent trend where pathways associated with amino acid metabolism were upregulated in cellulose-fed insects, while carbohydrate metabolism pathways were enriched by xylan. This effect was most notable in *Clostridiaceae* and *Oscillospiraceae 1,* which respectively upregulated eight and four carbohydrate pathways on the xylan diet, and eight and seven protein metabolism pathways when given cellulose (**Figure 3**). The remaining *Bacillota* had the same strong relationship between pathway and fiber with a few rare exceptions, such as cellulose enriching for C5-branched dibasic acid metabolism in *Lachnospiraceae C* and *Enterococcaceae* (**Figure 3A)**. Of all pangenomes, only *Desulfovibrio* showed exclusive enrichment by one fiber, with all significant metabolic pathways (C5-branched dibasic acid, lysine, tryptophan, arginine, and proline degradation) upregulated by cellulose.

**Figure 3:**
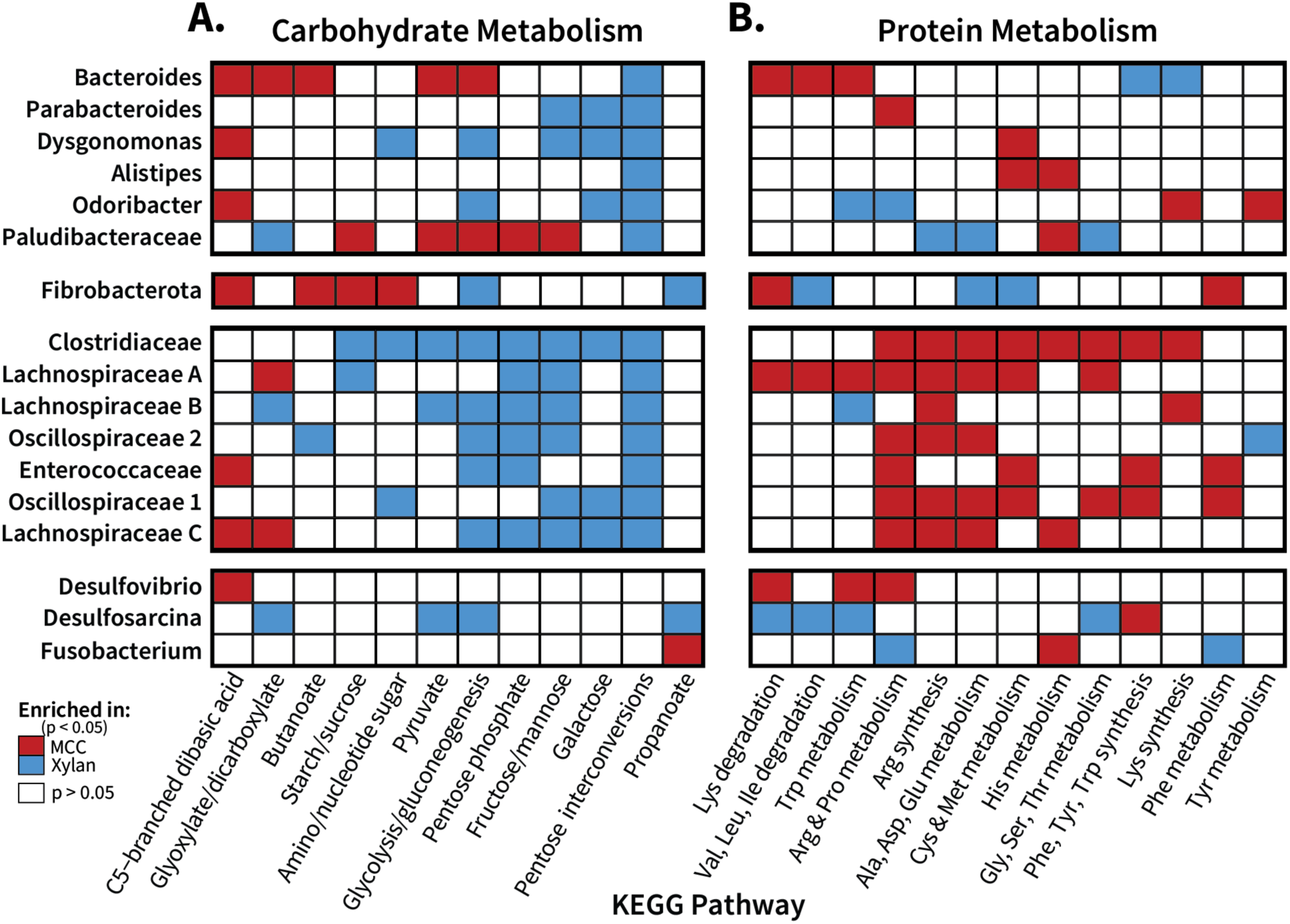
Pangenome organisms display fiber-dependent shifts in major metabolic pathways. Enrichment of (**A**) carbohydrate and (**B**) protein metabolic KEGG pathways was assessed for the pangenome transcriptomes of 100% xylan- and 100% MCC-fed insects. Gene clusters assigned relevant KOs were aggregated together based on pathway membership, which was then analyzed with DESeq2 at the pathway level to determine functional enrichment (adjusted p < 0.05) between the two fiber sources. All orthologs listed for the individual pathways were included into the overall pathway count, even if they appeared in multiple metabolic pathways.

In addition to pairwise comparisons between the single-fiber diets, we also explored changes in transcriptional activity for each pangenome across the ratio diets. Pangenome gene expression per ratio diets of KEGG pathways associated with carbohydrate metabolism and amino acid metabolism are displayed in **Supplemental Figures S7** and **S8,** respectively. Here, heatmap color indicates the relative abundance of transcripts assigned to each pathway within the individual pangenome’s transcriptome, while size indicates relative abundance of these transcripts out of all transcripts per sample/diet.

Across the ratios, *Bacteroides* transcripts associated with both carbohydrate (**Supplemental Figure S7A**) and protein (**Supplemental Figure S8A**) increased in overall abundance with increasing xylan content, consistent with the increase in total reads assigned to this organisms (**Figure 2).** However, when expressed in terms of relative abundance within the pangenome, *Bacteroides* transcribed more carbohydrate-associated genes in the cellulose diet, with the proportion of transcripts decreasing as xylan percent increased. Only pentose interconversion genes were both upregulated by *Bacteroides* on the xylan diet while also increasing in total abundance as dietary xylan percentage increased (**Figure 3**). Other *Bacteroidota* members showed less linear shifts in the overall abundance of carbohydrate metabolic pathways (**Supplemental Figure S7A**); of these, *Dysgonomonas* overall activity shared the positive correlation with xylan percent observed in *Bacteroides*, but differed in enrichment patterns. *Dysgonomonas* enriched for five pathways on xylan (nucleotide sugar; glycolysis; fructose/mannose; galactose; pentose interconversions) and transcribed its only cellulose-enriched pathway (C5-dibasic acid) at low levels both overall and within its transcriptome in all diet treatments. *Parabacteroides* and *Alistipes* maintained similar activity levels across all diets, although *Parabacteroides* did enrich for three sugar-related pathways on the xylan diet while *Alistipes* enriched only for pentose conversions. In contrast, *Odoribacter* and *Paludibacteraceae* displayed their highest activity levels on 100% cellulose, which dropped sharply once xylan percent reached 50%. *Paludibacteraceae* concurrently downregulated its expression in five pathways as xylan percent increased, matching its overall abundance shifts.

Amino acid metabolism pathways were generally transcribed at lower levels than the carbohydrate pathways among *Bacteroidota*, with less variation in expression observed over the diet gradient for most of the pangenomes (**Supplemental Figure S8A**). *Bacteroides* tryptophan metabolism showed a distinct switch from degradation on the 100% and 86% cellulose diet to synthesis on the 50% mix and xylan-majority diets, with a similar pattern in lysine metabolism. Among the other *Bacteroiodota* pangenomes, *Odoribacter* and *Paludibacteraceae* differentially expressed four pathways each, but generally amino acid pathway transcription remained consistent across the diet gradient.

Pangenomes within *Bacillota* exhibited both the highest overall activity and expression of carbohydrate pathway genes in mixed diets rather than either pure diet, particularly *Clostridiaceae, Lachnospiraceae,* and (to a lesser degree) *Enterococcaceae* (**Supplemental Figure S7B**). Carbohydrate metabolism in this group was largely enriched by xylan, but they displayed a clear preference for a mix of fibers over the 100% xylan diet. The *Oscillospiraceae* groups shifted their relative transcriptional activity based on diet, with carbohydrate pathways only enriched by xylan, but their absolute abundance of these pathways remained stable across the ratio diets with the exception of pentose interconversions and fructose/mannose metabolism in *Oscillospiraceae* group 1. Interestingly, while *Enterococcaceae* followed the general *Bacillota* trend of higher transcriptional activity on mixed diets, the opposite was true for galactose metabolism, where it had higher overall and relative transcription levels on both single fibers than it did on any mixed diet. Potentially, this group targets more host peritrophic matrix glycans during those diets while it is supported by a larger pool of intermediates during the *Bacillota* bloom on mixed diets.

*Bacillota* pangenomes showed similar patterns of overall transcript abundance for pathways associated with amino acid metabolism as they did carbohydrate metabolism, with the mixed diets resulting in higher overall activity levels for *Clostridiaceae* and *Lachnospiraceae* (**Supplemental Figure S8B)**. However, in terms of relative abundance within the pangenome, they transcribed pathways associated with protein metabolism at higher levels on the cellulose diet than any of the xylan containing diets. This trend is particularly marked for arginine and proline metabolism genes, which declined significantly as xylan was added to the diet for all *Bacillota* excluding *Lachnospiraceae B*.

Of the other phyla investigated, *Fibrobacterota* showed the clearest relationship between overall transcript abundance and cellulose in pathway expression of both carbohydrate (**Supplemental Figure S7C**) and amino acid metabolism (**Supplemental Figure S8C**). *Desulfobacterota* showed similar preferences for cellulose, although their expression levels decreased more gradually and remained higher in total over the gradient if diets than *Fibrobacterota*. *Fusobacterium* showed minimal alterations in either absolute abundance or relative transcriptional activity. In general, pangenomes showed phylum-distinct correlations with diet in carbohydrate and amino acid metabolism, indicating that *Bacillota* benefit more from a mixed-fiber environment while *Bacteroidota* possess a more linear relationship with the proportion of their choice fiber source.

### 3.5 Expression of genes for carbohydrate active enzymes

To evaluate the carbohydrate-degrading capabilities of the cockroach microbiome, we evaluated CAZyme diversity and abundance across the pangenomes. Overall, cockroach gut microbiota expressed an array of CAZyme families, with each pangenome encoding between 43 (*Fusobacterium*) and 255 (*Clostridiaceae*) unique CAZyme families (**Table 1)**. Glycoside hydrolases (GH) comprised the majority of CAZyme classes identified, followed by glycosyl transferases and polysaccharide lyases (**Supplemental Figure S9A).** In general, the diversity of CAZymes expressed did not correlate with transcript abundance, either within the metatranscriptome or as a fraction of transcripts within a given pangenome. By far, *Bacteroides* produced the greatest overall abundance of CAZyme transcripts, followed by the Bacteroidota groups *Parabacteroides* and *Dysgonomonas.* In contrast, while *Clostridiaceae* and *Lachnospiraceae B* (*Bacillota*) encoded the largest number of CAZyme families (**Supplemental Figure S9A)** and CAZyme-annotated gene clusters (**Supplemental Figure S9B)**, they produced fewer CAZyme transcripts, both in total and as a proportion of their transcriptional activity **(Figure 4A**, **Table 1)**. However, not all *Bacteroidota* expressed CAZymes at a high level; CAZymes made up <5% of total transcriptional activity among *Alistipes, Odoribacter* ,and *Paludibacteraceae* **(Table 1)**. While all *Bacteroidota* and nearly all *Bacillota* transcribed more glycoside hydrolases than any other class, the *Desulfobacterota* pangenomes and *Fusobacterium* instead produced higher levels of glycosyl transferases (**Figure 4A**), consistent with a focus instead on carbohydrate modification rather than degradation. *Oscillospiraceae* 2 deviated from the *Bacillota*, transcribing S-layer homology domains at higher levels than glycoside hydrolases. While not directly carbohydrate active, these proteins are often associated in *Bacillota* with CAZyme attachment to the cell wall.

**Figure 4:**
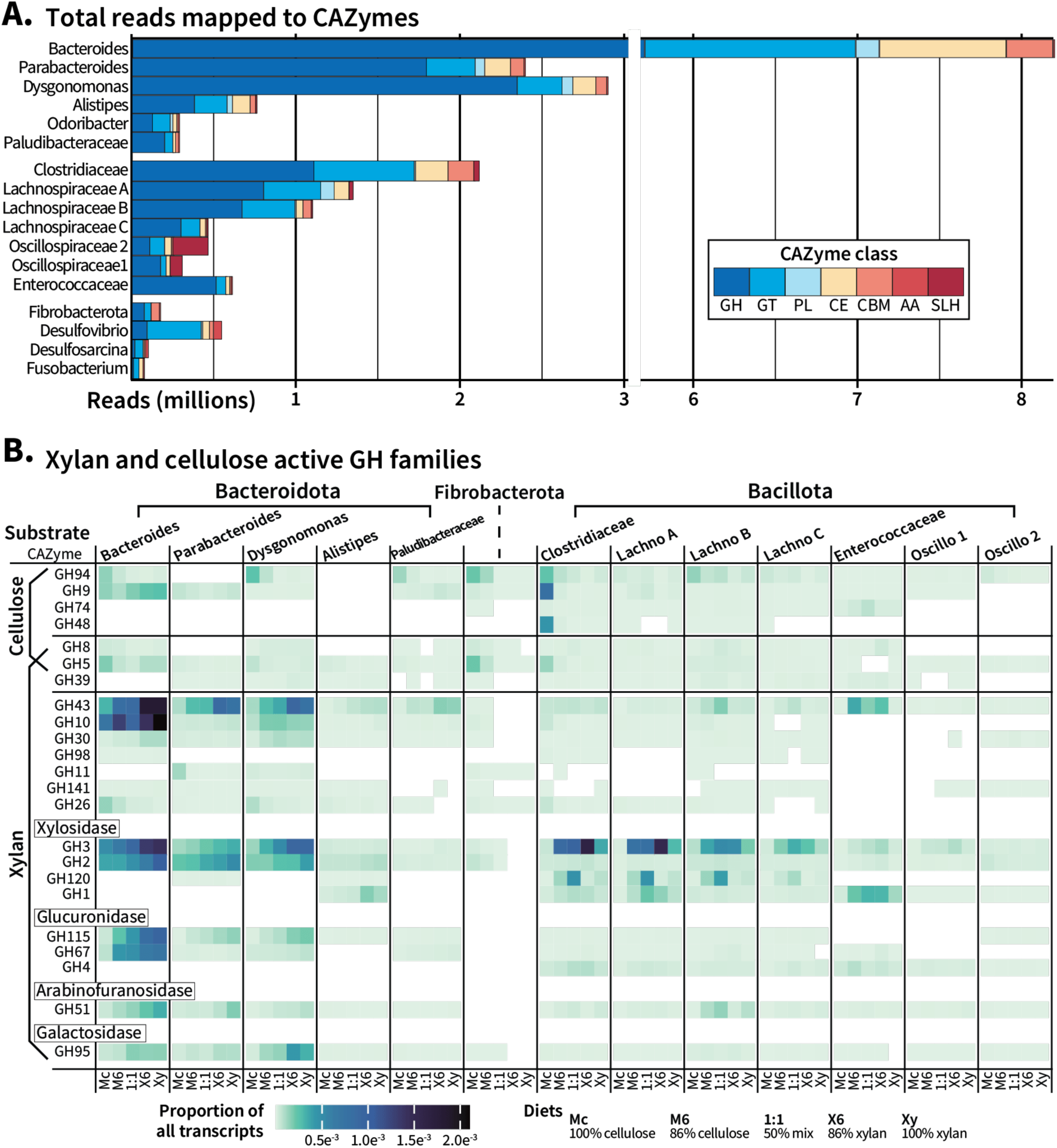
Glycoside hydrolase expression in pangenomes across the spectrum of fiber ratios. **(A)** The absolute abundance of CAZyme classes in each pangenome was calculated based on gene cluster transcripts (x 1,000,000) annotated within each class. **(B)** Gene clusters assigned to glycoside hydrolases (GH) associated with cellulose or xylan were aggregated and visualized with a heatmap scaled by proportion of these GH families out of all transcripts within the diets. Mc: 100% microcrystalline cellulose (MCC) diet; M6: 86% MCC diet; 1:1: 50% MCC/xylan diet; Xy: 100% xylan diet; X6: 86% xylan diet; Lachno: *Lachnospiraceae*; Oscillo: *Oscillospiraceae;* GH: glycoside hydrolase; GT: glycosyl transferase; PL: polysaccharide lyase; CE: carbohydrate esterase; AA: auxiliary activity; CBM: carbohydrate binding module; SLH: S-layer homology domain

To determine how the pangenomes responded to cellulose, xylan, and mixtures of the two as the primary dietary carbon source, we focused on CAZymes known to target these fibers, with special attention given to GH expression (**Figure 4B)**. We identified GH families in the pangenomes that are were identified as cellulases (GH94, GH9, GH74, GH48), xylanases (GH98, GH43, GH30, GH11, GH10), or contained activities for both (GH8, GH5, GH39) according to a substrate specificity file (**Supplemental Table 4**) downloaded from the dbCAN2 server [67].

Of the cellulases, GH94 was highly transcribed by members of *Bacteroidota* (*Bacteroides, Dysgonomonas, Paludibacteraceae*), *Fibrobacterota,* and *Bacillota* (*Clostridiaceae, Lachnospiraceae B)* on the 100% cellulose diet, with expression levels decreasing as the dietary cellulose:xylan ratio decreased. GH9 was highly expressed in *Bacteroides,* but did not seem to consistently correlate with dietary cellulose levels in the *Bacteroidota;* only *Clostridiaceae* produced more of this enzyme in the cellulose diet. While GH8, 5, and 39 were classified as processing both fibers, only GH5 expression differed across the diets. GH5 was transcribed heavily by *Fibrobacterota* and *Clostridiaceae* on the cellulose diet, with *Bacteroides* expressing gene clusters with this CAZyme at both the 100% cellulose and xylan diets; the GH5 family contains several subfamilies with targets including arabinan, chitin, and β-mannan that may be the source of this variation (**Supplemental Table 4)**.

Since xylan is a complex and heterogenous polysaccharide with wide variance in the side chain composition depending on its source, we divided xylan-associated glycoside hydrolases based on their most probable targets (**Figure 4B)**. Breakdown of xylan requires the successive and cooperative activity of glycosidases that remove sugar residues, endo-xylanases that cleave inner xylose-xylose bonds, and exo-xylanases or xylosidases that remove terminal xylose or xylooligosaccharides.

CAZyme families associated with side chain removal include hydrolases targeting glucuronic acid (GH115, GH67, GH4), arabinose (GH51), and galactose (GH95) side chains. Most of these enzymes were produced at high levels by *Bacteroides, Parabacteroides,* and *Dysgonomonas,* with the exception of GH4, which was absent in all *Bacteroidota.* Of the two glucuronidases present in this phylum, enzymes from family GH115 increased for these three pangenomes as xylan increased, while GH67 expression only responded to dietary xylan in *Bacteroides.* Of the other sugars, *Dysgonomonas* transcribed the most galactosidase (GH95), which was positively correlated with dietary xylan, but only marginally increased in arabinofuranosidase activity. Rather, *Parabacteroides* and *Bacteroides* produced steeper diet-associated gradients of GH51 family enzyme expression. In contrast, *Bacillota* pangenomes in general did not express CAZymes associated with hydrolyzation of these side chains at high levels or in response to dietary xylan. *Lachnospiraceae* B upregulated GH51 family enzymes in the mixed diets while *Clostridiaceae* and *Enterococcaceae* increased GH4 family enzymes targeting glucuronic acid, but the shifts in transcription for these enzymes were subtle and their expression was generally low in *Bacillota* pangenomes independent of diet.

Endo-xylanases are found in CAZyme families GH141, GH98, GH43, GH30, GH26, GH11, and GH10. Of these, GH43 stands out with the strongest correlations to dietary xylan percent, with all *Bacteroidota* members showing gradual increases in expression as percent xylan increased. In *Bacteroides,* GH10 shows similar upregulation correlated with diet, while *Dysgonomonas* appears to maintain more sustained levels once xylan is added to the diet in any amount. Curiously, GH11 and GH26 presented opposite patterns than expected, with higher levels found on the cellulose diet that suggest these enzymes are targeting glucans rather than xylan. In *Bacillota*, GH43 expression is increased on the mixed diets in *Enterococcaceae*, *Lachnospiraceae B*, and *Lachnospiraceae C,* but otherwise xylanases in this phylum are expressed at low levels, if at all.

Finally, xylosidases are associated with the removal of terminal xylose during breakdown of shorter, lower complexity xylo-oligosaccharides. *Bacteroidota* primarily expressed xylosidases from families GH3 and GH2 in the characteristic gradient-like fashion, with *Dysgonomonas* producing more xylosidases than they did xylanases. *Bacteroides* and *Parabacteroides* produced high levels of GH3 xylosidases but expression of these enzymes remained below their GH43 xylanase expression. *Bacillota,* unlike with xylanases, did express GH3 xylosidases at high levels and significantly upregulated their expression in response to increasing dietary xylan content, which dropped once cellulose was absent in the diet. In addition to GH3, *Clostridiaceae* and *Lachnospiraceae* produced GH120 xylosidases, which consistently peaked in expression at the 50% mixed diet then dropped once xylan was the majority fiber. Overall, based on CAZyme profiles obtained for these pangenomes, *Bacteroidota* in the cockroach gut microbiome are responsible for primary xylan deconstruction as well as the majority of side chain removal while *Bacillota* liberate terminal xylose residues.

### 3.6 Phylum level differences in core carbon metabolic activity

To decipher the specific contributions of cockroach gut microbiota to xylan and cellulose degradation and how these different groups utilized the metabolic products, we compared phylum-level expression of KEGG orthologs catalyzing steps involved in carbohydrate catabolism and central carbon metabolism (**Figure 5**). The KEGG modules for glycolysis (M00001), pentose phosphate pathway (M00004), and the Entner-Doudoroff shunt (M00308) were incorporated into a single reaction map with curated KOs related to glucuronic-arabinoxylan and microcrystalline cellulose degradation. For KEGG ortholog IDs and their fiber-based enrichment for the individual pangenomes, please see **Supplemental Figure S10**.

**Figure 5:**
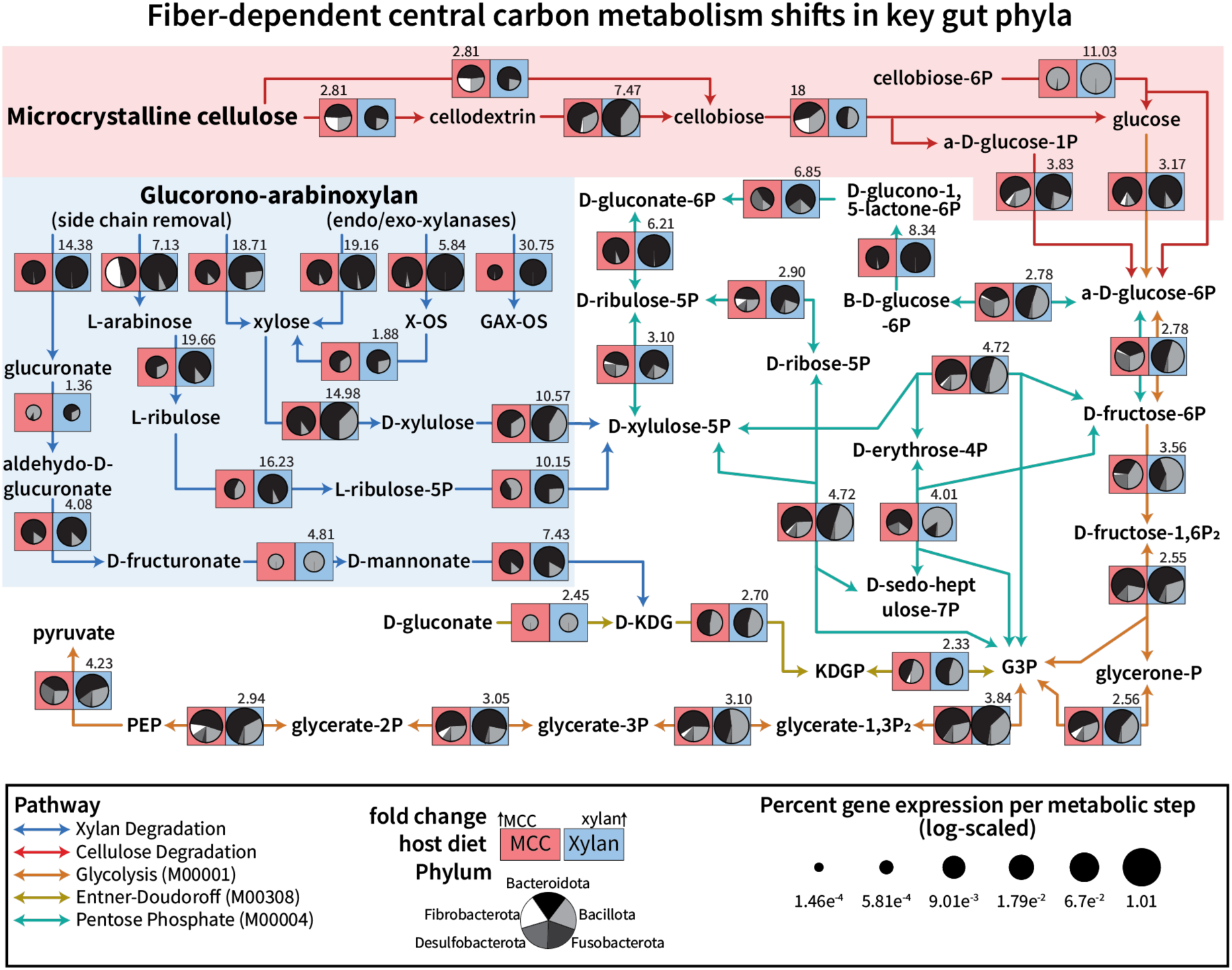
Key cockroach phyla demonstrate flexibility in central carbon metabolism when shifting between degradation of xylan and cellulose. Pangenome gene clusters containing KEGG orthologs involved in glycolysis, the Entner-Doudoroff pathway, and the pentose phosphate pathway, as well as curated KOs involved in xylan and cellulose degradation (see **Supplemental** Figure 10), were converted based on enzymatic step to their relative abundance of total transcripts and averaged by diet. Enzymatic steps within the pangenomes were summed together by phylum, and the total contribution of *Bacteroidota*, *Bacillota*, *Desulfobacterota, Fusobacterota,* and *Fibrobacterota* to each step was visualized with pie charts. The size of the pie charts indicates the overall transcriptional percent that step comprises in the 100% cellulose (red box) and 100% xylan (blue box) diets, with the number indicating the fold increase in relative abundance in the direction of either cellulose (left corner) or xylan (right corner). Pathways are indicated with arrow color, and the transparent boxes are included to focus attention on xylan degradation steps (blue) and cellulose degradation steps (red). MCC = microcrystalline cellulose; G3P = glyceraldehyde-3-phosphate; PEP = phosphoenolpyruvate

Cellulose is associated with significant increases in few KEGG orthologs, which can be split into three main activity types: endoglucanase (K01179), beta-glucosidase (K05349, K05350), and cellobiose phosphorylase (K00702). Endoglucanase activity, which produces cellodextrins or cellobiose from crystalline cellulose, was higher in the cellulose diet than xylan with much of the increase driven by *Fibrobacterota* activity, although several *Bacteroidota* and *Bacillota* pangenomes did show cellulose-based enrichment as well on this step. Cellobiose phosphorylase activity increased 18 fold on the cellulose diet, again driven by *Fibrobacterota* and supported by enrichment of fiber-degrading pangenomes. Orthologs for beta-glucosidases that may cleave cellodextrins into a glucose plus cellobiose were included, but only *Fibrobacterota* and *Oscillospiraceae* group 2 enriched for one of these KOs on cellulose; K05349 was enriched by xylan in six *Bacillota* organisms and three *Bacteroidota*, explaining the larger portion of transcripts found in the hemicellulose diet compared to cellulose (**Supplemental Figure S10**). Glycolysis genes were higher in the xylan diet than cellulose, but this may be due to the high activity and abundance of *Bacteroides* biasing the overall numbers. Notably, *Desulfobacterota* glycolytic activity was correlated strongly with the cellulose diet, but neither of those pangenomes could degrade cellulose or cellobiose on their own.

During xylan degradation, endo-xylanase activity was almost exclusively performed by *Bacteroidota* (K01181, K15924, K01198), while *Bacillota* showed a large proportion of xylosidase activity (K01811), consistent with the CAZyme profiles presented in **Figure 4**. *Bacillota* then converted xylose to D-xylulose (xylose isomerase; K01805) and phosphorylated the intermediate (xylulokinase; K00854) for funneling into the pentose phosphate pathway for central carbon metabolism. *Bacteroidota* did cleave xylose residues off to feed into central metabolism as well, but additionally demonstrated increased arabinose (K01209, K15921) and glucoronate (K01235) removal, with the arabinose feeding into the pentose phosphate pathway through conversion to L-ribulose (K01804), L-ribulose-5P (K00853), and finally D-xylulose-5P (K03077). *Bacillota* pangenomes did enrich for arabinose cleavage and conversion (**Supplemental Figure S10**), but to a lesser degree than *Bacteroidota* and less so than they did xylose. Once D-xylulose-5P is obtained, these two phyla demonstrated different strategies; *Bacteroidota* increased their transcription of enzymes guiding the pentose into glycolysis as glucose, while *Bacillota* seemed to prefer transcribing higher pentose phosphate pathway genes that lead to intermediary glyceraldehyde-3-phosphate.

None of the pangenomes encoded a complete Entner-Doudoroff shunt, but glucuronate removed from glucuronoxylan could be converted to 2-keto-3-deoxy-gluconate (KDG) to form the intermediate glyceraldehyde-3-phosphate (G3P). While glucuronate conversion steps were inconsistently expressed by these pangenomes, both *Bacillota* and *Bacteroidota* contribute similar proportions of transcripts to this shunt and increased their transcription on the xylan diet, suggesting they may leverage this pathway as needed. Taken together, these results shed light on the strategies leveraged by a complex community of gut microbiota to degrade dietary fibers and incorporate the component sugars into their central carbon metabolism.

## 4 Discussion

Dietary fiber is an important driver of gut microbiome structure, and diets containing purified fibers enable the individual contributions of microbial community members to polysaccharide degradation to be teased apart even within complex gut systems. This work sought to identify the mechanisms behind large-scale compositional shifts observed previously in cockroach gut microbiota when exposed to synthetic diets containing purified fibers as the sole carbohydrate source [21]. To do this, we fed cockroaches synthetic diets containing as the carbohydrate source xylan, microcrystalline cellulose, or a mixture of both polysaccharides in differing ratios to detect fiber-dependent microbial responses and determine how mixed fibers influence these dynamics. We tested the impact of these diets on the hindgut microbiome composition using 16S rRNA gene amplicon sequencing, followed by functional characterization of the microbiota through metatranscriptomic analysis. Leveraging an organism-centric pangenome approach, we characterized the metabolic strategies enriched in these organisms by the individual fibers and discovered phylum-dependent responses to the different ratios of xylan:cellulose in mixed fiber diets.

### Compositional Overview

As previously observed [21], the hindgut microbiota from cockroaches fed xylan or cellulose differ in diversity and taxonomic composition of 16S rRNA profiles; xylan-fed communities are less diverse across multiple measures than cellulose-fed communities (**Figure 1**). Expanding on these results using mixed-fiber diets, we found that diets containing mixtures of dietary xylan and cellulose select for a gradient in gut microbiome composition that reflects the influence of both fibers (**Figure 1**). However, the relationship between the dietary xylan:cellulose ratio and gut community composition was not linear. Mixed diets showed substantially higher alpha diversity than the 100% xylan regardless of their own xylan content (**Figure 1A-C**). In contrast, xylan content strongly influenced beta diversity measures, particularly those that consider taxon abundance, with clear separation of 100% cellulose diets from even low-xylan mixed diets (**Figure 1D-E**).

We compared the composition of our 16S rRNA results those of our metatranscriptomes (**Figure 2A, S3**), which we calculated based on top-scoring taxonomic hits obtained from read alignment to both the nonredundant RefSeq protein database and cockroach-associated single cell genomes [22]. Alpha and beta diversity of metatranscriptome compositions reflected the core differences observed between the 100% xylan and 100% cellulose diets, highlighting that xylan generates a functionally distinct, less diverse range of active microbes compared to cellulose (**Supplemental Figure S4**).

Interestingly, the mixed-fiber metatranscriptomes seemed to have stronger majority-fiber affiliations in their alpha and beta diversity outcomes than they did in the amplicon dataset, while the 50% mix produced the widest sample variation (**Supplemental Figure S4**). Whether this variability is an artifact of our samples or due to destabilizing effects of the fiber source is difficult to say; literature on the functional implications of mixing purified fiber sources in the complex microbiome of a living host is limited [72], while *in vitro* studies produce divergent results. Previous work modeling responses of human donor microbiota found in one case that individual fiber complexity was more impactful than mixing multiple (3 or 6) fiber sources [73] and in another, that multiple fibers increased diversity more so than their component fibers [74].

While the taxonomic compositions of these two datasets were generally consistent, we did observe some differences in their relative abundance of major bacterial orders (**Figure 2A**), partially due to the different taxonomies utilized by the different data processing methods (SILVA taxonomy for 16S rRNA gene amplicons vs NCBI taxonomy for metatranscriptomes) in the case of *Christensenellales* and *Oscillospirales*. Other differences came from *Lactobacillales,* which were identified as abundant and enriched by xylan in the 16S dataset but not the metatranscriptome, and in *Bacteroidales,* which displayed substantially higher activity in the metatranscriptome than expected based on 16S rRNA gene abundance. While the large increase in *Bacteroidales* abundance in the metatranscriptomes likely indicates that this group is highly transcriptionally active, it is unclear whether the diminished *Lactobacillales* population reflects a real population with low transcriptional activity, or if there are differences in the taxonomic assignment of transcripts vs 16S rRNA amplicons. Similar loss of *Lactobacillaceae* in metagenome results despite apparent 16S rRNA presence has been reported in the cockroach [22], suggesting that amplicon sequencing may inaccurately capture this group of organisms.

### Pangenome Activity

A major challenge in metatranscriptome analyses lies in transcript mapping and reference genome selection. This is a particular challenge for less-studied models such as the cockroach, where relatively few gut taxa have been cultured and sequenced. To overcome this challenge, we used an approach that mapped transcripts to pangenomes constructed from cockroach-originated single cell genomes [22] and closely related reference genomes from other environments. [56, 70]. In all diets, most reads were successfully assigned to one of 17 pangenomes of cockroach gut symbionts (**Figure 2B**), allowing us to analyze the fiber-dependent functional behavior of these organisms via their individual “transcriptomes”, as well as identify phylum-wide patterns when faced with a gradient of fibers.

Further, this approach allows us to discern true differences that arise in the microbiota due to dietary fiber type. While the frequency of ASVs enriched per each genus may suggest a preference towards xylan or cellulose, abundant ASVs enriched by the two different fibers in the 16S rRNA dataset notably overlap in in taxonomic origin (**Supplemental Figure 3**). Major organisms with ASVs enriched by both fibers include Bacteroidota members *Bacteroides, Dysgonomonadaceae,* and *Parabacteroides,* while *Alistipes* and *Odoribacter* had members enriched by xylan or the mixed fibers, all of which were investigated using our pangenome approach and discussed below.

*Odoribacter* in particular shows increased activity on cellulose in metatranscriptomic analysis despite its enrichment by xylan in the 16S data set. Enriched *Bacillota* in the 16S data on both fiber sources include *Lachnospiraceae* members, *Ruminococcus*, and *Christensenellaceae,* although many more members were enriched by xylan than cellulose. *Ruminococcus* and *Christensenellaceae* are included in the *Oscillospiraceae* pangenome groups, but ended up being less important in *Bacillota* activity; rather, *Clostridiaceae*, which were not represented in the 16S, are the dominant *Bacillota* group in the metatranscriptome. *Desulfobacterota* and *Fibrobacterota* were enriched by cellulose in both 16S abundance and in transcriptomic activity. Through the pangenomes, we are able to confirm the dynamics suggested by 16S rRNA gene abundance, while clarifying apparent discrepancies that arose, highlighting the value of this approach.

#### Bacteroidota

*Bacteroidota* are among the most abundant organisms found in omnivorous gut communities, including omnivorous cockroaches [20, 21, 23, 25, 75]. Here, we analyzed six *Bacteroidota* pangenomes found abundantly in the cockroach hindgut: *Bacteroides, Parabacteroides, Dysgonomonas, Alistipes, Odoribacter,* and *Paludibacteraceae.* Among these organisms, we found *Bacteroides* to be especially active regardless of dietary fiber source (**Figure 2B**, **Table 1**), consistent with their proposed role as major fiber-degrading players in the cockroach gut [22, 23]. In human gut systems, *Bacteroides* are particularly well-known for their ability to break down an array of fibers including xylan [76, 77], starch [78], host glycans [79, 80], cellulose [23], and pectin [81] due to their large assortment of encoded carbohydrate-active enzymes (CAZymes) and the use of polysaccharide utilization loci (PULs) to maximize substrate compatibility. We observed the same CAZyme diversity and expression here, with *Bacteroides* transcribing 175 unique CAZyme families (**Supplemental Figure S9A**) at very high levels (**Figure 4A**), vastly outnumbering CAZyme transcription by all other organisms. Two other *Bacteroidota* groups, *Dysgonomonas* and *Parabacteroides,* were the second and third largest source of CAZyme transcripts, consistent with a role as primary fiber degraders.

*Bacteroides, Dysgonomonas,* and *Parabacteroides* transcriptional profiles suggest a major role in metabolism of dietary xylan. These three *Bacteroidota* groups expressed CAZyme families that are active on common xylan branches, including orthologs responsible for cleaving and subsequently processing arabinose residues, and also displayed primary chain xylanase activity, which together argue towards their ability to utilize complex xylan structures (**Figure 4B**).

Ordination analysis of transcriptional profiles from all three groups showed a strong response to diet (**Supplemental Figure S5A**), including a strong significant response to dietary xylan percentage and, to a lesser extent, presence/absence of xylan, but not presence/absence of cellulose. This, together with their high expression of xylan degradation genes and much weaker expression of cellulolytic genes suggests a strong preference for xylan rather than cellulose as a growth substrate. Given these results, we hypothesize that these groups are the primary xylan degraders in the cockroach gut. The persistence of these organisms in cockroaches fed a pure cellulose diet plus their enrichment of cellulase and cellobiose phosphorylase transcripts suggest that they may be able to metabolize cellulose if necessary. However, inferring this definitively is difficult given their persistence during long-term host starvation [20] and their ability to leverage host glycans for sustenance [79].

#### Bacillota

While *Bacillota* are commonly identified in omnivorous gut communities, they are usually classified as secondary fermenters rather than primary degraders [22, 75, 82]. Our results are consistent with a role as secondary fermenters of xylan breakdown products; *Bacillota* organisms were generally associated with dietary xylan content in their 16S rRNA gene abundance with more enriched ASVs than in cellulose (**Supplemental Figures S2, S3**), and pangenome transcriptional activity enriched overall by xylan presence (**Figure 2, Supplemental Figure S7, S8**) but primarily produced xylosidases rather than endo-xylanases or glycosidases active on side chains (**Figure 4B**). This was correlated with xylan-enriched transcription of xylose isomerase, xylulokinase, and pentose phosphate pathway orthologs feeding into G3P production (**Figure 5, S10**), illustrating the xylose-utilizing potential of Firmicutes [83, 84]. Interestingly, *Bacillota* transcriptional profiles were better explained by models considering absence/presence of xylan rather than dietary xylan percentage, consistent with a large metabolic response to dietary xylan at even the lowest abundance tested.

*Bacillota* displayed clear preferences for xylan as a carbohydrate source, upregulating between 3-8 carbohydrate degradation pathways per organism, while on the cellulose diet they enriched instead for numerous amino acid metabolic pathways (**Figure 3),** suggesting a switch to utilization of dietary amino acids/proteins in cellulose diets, rather than secondary fermentation of cellulosic substrates.

There are cellulose-active *Bacillota* found in herbivorous communities, such as *Ruminococcus* species [85, 86] or cellulosome-producing *Clostridium* [87], but the *Clostridiaceae* here showed higher preference for xylan (**Figure 2B, Supplemental Figure S7, S8**), and while *Oscillospiraceae* 2 was slightly more abundant on cellulose than xylan (**Supplemental Figure S7**) it shared the KEGG pathway enrichment patterns observed in the other *Bacillota* pangenomes (**Figure 3**). Generally, *Bacillota* pangenomes had relatively similar metabolic activity allocations across the different xylan-containing diets, but surprisingly, displayed distinctly higher overall transcriptional activity in mixed-fiber diets (**Supplemental Figure S7, S8**). Although the apparent switch to amino acid-degrading vs. carbohydrate metabolic genes in cellulose fed cockroaches suggests that they may not be as actively involved in cellulose fermentation as xylan, they may still benefit from intermediates being released by cellulose-degrading organisms, either as catabolic substrates or as anabolic precursors for biomass production [88].

#### Fibrobacterota

Unexpectedly, we discovered a significant increase in *Fibrobacterota* transcripts in the 100% cellulose diet. This small phylum contains well-characterized members of ruminant and herbivore monogastric gut communities that are rarely identified or studied in omnivorous animals [89, 90], although we did identify a small presence of this group in our previous 16S rRNA amplicon survey of *P. americana* [21]. *Fibrobacterota* are remarkably specialized towards cellulose metabolism, encoding a unique suite of genes that facilitate fiber degradation by releasing cellulases and hemicellulases into the surrounding area prior to transporting cellodextrins into the cell for degradation [91–93]. In line with this, we did observe cellulose-associated CAZyme transcription (**Figure 4B**), in addition to high expression of cellulase ortholog transcription in the cellulose diet (**Figure 5**). Of the pangenomes assessed, *Fibrobacterota* were one of only 2 organisms where cellulose presence/absence significantly impacted overall transcriptional profile of that organism (**Table 2**), and unconstrained ordinations showed strong separation of transcriptional activity under high-cellulose (86% and 100%) diets. The sharp shift in transcriptional activity in diets with 50% or greater xylan content suggest that gut microbial xylan metabolism may interfere with cellulose degradation by Fibrobacterota. These organisms are known to be highly sensitive to catabolite repression, and environmental pH [94]; the bloom in *Bacillota,* for example, likely altered altered gut pH or metabolite profiles [95, 96].

When comparing pangenome activity patterns in carbohydrate metabolic pathways across diet ratios (**Supplemental Figure S7**), we noticed that four additional organisms displayed their highest activity on cellulose: the *Desulfobacterota* pangenomes *Desulfosarcina* and *Desulfovibrio*, and *Bacteroidota* members *Paludibacteraceae* and *Odoribacter*. With the exception of *Desulfovibrio*, these were among the lower activity organisms in our metatranscriptomes, with little fiber-degrading activity themselves (**Table 1**: <5% of expressed gene clusters labeled as CAZymes), leading us to consider that *Fibrobacterota* may be supporting their higher growth in this diet. One characteristic of *Fibrobacterota* in rumen and gut communities is its role in cross-feeding; these organisms commonly provide neighbors with cellobiose and hemicellulose-derived oligo- and monosaccharides produced by their activity for the sole purpose of increasing cellulose accessibility, and they release metabolic waste products succinate and formate [97–100]. We observed that transcription of succinate dehydrogenase KOs were enriched in *Fibrobacterota* from the cellulose diet compared to the xylan diet (K00241, K00240, K00239; **Supplemental Figure S10, S11**), suggesting a similar function in cockroach-derived species. *Desulfobacterota* have previously been established as hydrogen and formate cross-feeders associated with cellulose degrading communities, strengthening the possibility of a relationship with cellulose-degrading *Fibrobacterota* [101–103]. *Paludibacteraceae* expressed higher GH94 family CAZymes on the cellulose diet, which typically has cellobiose phosphorylase activity (**Figure 4D**) and this organism showed similar substantial drops in carbohydrate activity as seen in *Fibrobacterota* as cellulose percent decreased (**Supplemental Figure S7**). This suggests that *Paludibacteraceae* may benefit from the cellobiose liberated by *Fibrobacterota.* Finally, *Odoribacter* has not been previously reported to be an effective fiber degrader, but it has been hypothesized to consume succinate in mouse models [104, 105], and may be utilizing succinate produced by *Fibrobacterota*. While *Fibrobacterota* is an uncommon member of omnivore microbiomes, these findings suggest they may play similar roles in the cockroach gut as they do in herbivores, mediating cross-feeding interactions that increase and maintain diversity in complex gut systems.

## Conclusions

In conclusion, through 16S rRNA gene profiling combined with RNA-seq, we discovered that xylan and cellulose differentially influence the community structure and function of the American cockroach hindgut microbiome. A noteworthy observation from this study was that the composition predicted by 16S rRNA gene sequencing was not necessarily reflected in the transcriptional activity of the gut community. In particular, *Bacteroides* was far more active than estimated by its amplicon abundance, and their transcriptional profile was consistent with a role as primary degraders of xylan despite the fact that they declined in relative abundance as measured by 16S rRNA gene assays of xylan-fed cockroaches. In contrast, *Lachnospirales* exhibited a very strong, significant relationship with xylan as measured by ASV abundance, but transcriptional profiles suggest a secondary role as fermenters of xylosides. In addition, they displayed far higher transcriptional activity in the mixed-fiber diet than on pure xylan. These results stress the importance of combining functional analysis with amplicon surveys to fully grasp the dynamics occurring within a complex gut community.

Through our organism-centric pangenome approach, we identified *Bacteroidota* as likely to be responsible for most xylan degradation in the cockroach gut, with members *Bacteroides*, *Parabacteroides,* and *Dysgonomonas* producing high levels of diverse CAZymes that increased with dietary xylan percent. By mixing these fibers in different ratios, we discovered that *Bacillota* members were enriched by xylan but flourished in abundance and activity when given mixed fibers, benefiting from their own metabolic activity as well as the actions of other fiber-degrading organisms. Finally, we observed surprisingly high *Fibrobacterota* cellulolytic activity on the cellulose diet. *Fibrobacterota* is an uncommon organism in omnivorous gut microbiomes; the ability of the cockroach to host these organisms may contribute to their famous ability to survive long-term on nutrient-poor diets. These results underscore the power of metatranscriptomic approaches combined with synthetic diets to asses gut microbial responses to purified or mixed dietary fibers, and highlight the utility of the omnivorous cockroach as a model to better understand complex microbial interactions in the guts of omnivores.

## 5 Data Availability

Data associated with this study are available from the NCBI Sequence Read Archive. Raw 16S amplicon data can be found under BioProject PRJNA1105088 (SRA experiments: SRX24377840 SRX24377655 - SRX24377840).

