## Supplemental Figures and Tables for "Niche specialization and cross-feeding interactions shaping gut microbial fiber degradation in a model omnivore"

**Supplemental Tables and Figures**

**Supplemental Table 1: Primers for generating antisense rRNA probes.** DNA was extracted from two pooled cockroaches that had been fed synthetic diets containing either xylan or microcrystalline cellulose. Microbial and host-derived rRNA gene regions were amplified with the T7 promoter region appended to the reverse barcode using the primers and annealing temperatures listed in this table.

| Region | Primer | Sequence | Anneal  (C) |
| --- | --- | --- | --- |
| Archaea 16S | 21F | TCCGGTTGATCCYGCCGG | 70 |
|  | 1492R | GCCAGTGAATTG-**T7**-GGGGYYACCTTGTTACGACTT |  |
| Archaea 23S | 189F | ASAGGGTGAHARYCCCGTA | 70 |
|  | 2490R | GCCAGTGAATTG-**T7**-GGCTGTCTCRCGACGGTCTRAACCCA |  |
| Bacteria 16S | 27F | AGAGTTTGATCCTGGCTCAG | 39 |
|  | 1492R | GCCAGTGAATTG-**T7**-GGACGGCTACCTTGTTACGACTT |  |
| Bacteria 23S | 189F | GAASTGAAACATCTHAGTA | 39 |
|  | 2490R | GCCAGTGAATTG-**T7**-GGCGACATCGAGGTGCCAAAC |  |
| Eukaryote 18S | 1F | ACCTGGTTGATCCTGCCAG | 55 |
|  | 1520R | AATTA-**T7**-ATTCYGCAGGTTCACCTAC |  |
| Eukaryote 28S | 26F | ACCCGCYGAAYTTAAGCATA | 55 |
|  | 3126R | AATTA-**T7**-ATTCTGRYTTAGAGGCGTTCAG |  |
| Cockroach  ITS | 18S_1907F | CCTGCGGAAGGATCATTAAC | 60 |
|  | 28S_-2R | GCCAGTGAATTG-**T7**-GGCTTAAATTCAGCGGGTAGTCTC |  |

**T7 sequence**: 5′ TAATACGACTCACTATAG 3′

**Supplemental Table 2: RNA read tracking through filtering steps.**

|  | **R1 Reads** | | **R2 Reads** | | **Both R1 and R2** | | | |
| --- | --- | --- | --- | --- | --- | --- | --- | --- |
| **Sample** | **total** | **mRNA** | **total** | **mRNA** | **paired reads** | **filtered reads** | **merged pairs** | **aligned with DIAMOND** |
| **XT1_HgL1** | 83167708 | 25239059 | 83643619 | 25592007 | 49090264 | 45249236 | 22624618 | 18085988 |
| **XT1_HgL2** | 47251549 | 13717591 | 47902798 | 13999554 | 26871502 | 22956298 | 11478149 | 9256938 |
| **XT1_HgL3** | 79007875 | 25474355 | 79996235 | 25923374 | 49611730 | 38464086 | 19232043 | 15116932 |
| **XT1_HgL4** | 68303715 | 22841708 | 69863332 | 23089066 | 44978238 | 41017224 | 20508612 | 17093359 |
| **XT1_HgL5** | 62065220 | 25785807 | 62951191 | 25990256 | 50609950 | 48830212 | 24415106 | 20288972 |
| **XT1_HgL7** | 91950695 | 37851719 | 94496533 | 38063774 | 74326902 | 71794094 | 35897047 | 26883520 |
| **XT2_HgL1** | 71418991 | 25076230 | 72106314 | 25243676 | 48799680 | 47469354 | 23734677 | 19161622 |
| **XT2_HgL3** | 52469393 | 19524666 | 52759634 | 19687352 | 38619820 | 37445000 | 18722500 | 14284306 |
| **XT2_HgL4** | 83984458 | 39400506 | 84688749 | 39821140 | 77030924 | 75385452 | 37692726 | 31943483 |
| **XT2_HgL5** | 79415147 | 36860338 | 80928173 | 37043447 | 72835484 | 70430846 | 35215423 | 29295230 |
| **XT2_HgL6** | 67709119 | 29534363 | 69153831 | 29887045 | 57954520 | 55633270 | 27816635 | 23655953 |
| **XT2_HgL7** | 73025327 | 29103049 | 74984102 | 29950680 | 55444638 | 53052316 | 26526158 | 18876312 |
| **XT3_HgL1** | 46175011 | 16233127 | 49777036 | 16559143 | 31718522 | 30124226 | 15062113 | 14115915 |
| **XT3_HgL3** | 60384706 | 28281236 | 63055728 | 28448960 | 55805312 | 53821444 | 26910722 | 22514244 |
| **XT3_HgL4** | 89610533 | 30366146 | 90682727 | 30824303 | 59864898 | 54306166 | 27153083 | 23595116 |
| **XT3_HgL5** | 107781761 | 28649132 | 109234848 | 28848385 | 55647064 | 54225038 | 27112519 | 21560649 |
| **XT3_HgL9** | 73214318 | 11597279 | 73430946 | 12162830 | 22157234 | 16997708 | 8498854 | 7144408 |
| **XT3_HgL10** | 69422560 | 30198829 | 69952461 | 30316467 | 59390396 | 51683960 | 25841980 | 21365461 |
| **XT4_HgL6** | 54613882 | 13862109 | 54815672 | 14181923 | 25969036 | 24460134 | 12230067 | 8382119 |
| **XT4_HgL7** | 83302304 | 24855380 | 84860431 | 25080399 | 48623788 | 45523768 | 22761884 | 16206823 |
| **XT4_HgL8** | 75189344 | 16448440 | 75715652 | 16750454 | 31794802 | 28956358 | 14478179 | 10279389 |
| **XT4_HgL9** | 50606869 | 13376743 | 52100115 | 13695932 | 25613404 | 23274734 | 11637367 | 8419598 |
| **XT4_HgL10** | 102161190 | 32863172 | 102894236 | 33007141 | 64572796 | 62291220 | 31145610 | 23975956 |
| **XT4_HgL11** | 95711249 | 25753521 | 96394620 | 25776625 | 49656214 | 44609196 | 22304598 | 13918250 |
| **XT5_HgL3** | 65446044 | 18930397 | 66687129 | 19144507 | 37124530 | 34316272 | 17158136 | 13187727 |
| **XT5_HgL4** | 38712542 | 12923662 | 40478389 | 13043323 | 24976818 | 23244390 | 11622195 | 7978854 |
| **XT5_HgL5** | 81916125 | 7545610 | 82014303 | 7931598 | 13903986 | 13154186 | 6577093 | 4875506 |
| **XT5_HgL6** | 53985913 | 16666582 | 54300297 | 16780659 | 32513704 | 30386132 | 15193066 | 11545523 |
| **XT5_HgL7** | 115045888 | 8710681 | 115193918 | 9554599 | 15689626 | 12026460 | 6013230 | 4740617 |
| **XT5_HgL10** | 136881495 | 12508722 | 137154114 | 13039750 | 23307506 | 18195732 | 9097866 | 6273811 |

**Supplemental Table 3: RefSeq accessions used to generate *Lachnospiraceae* pangenome groups.**

| **Genera** | **Accessions** |
| --- | --- |
| ***Lachnospiraceae A*** | |
| *Aequitasia* | GCF_024160205\|GCF_024721305\|GCF_024160185\|GCF_024721265 |
| *Brotaphodocola* | GCF_020686985\|GCF_003478505\|GCF_003477935\|GCF_003480315\|GCF_003480105\|GCF_003481985\|GCF_003481825 |
| *Enterocloster* | GCF_000158075\|GCF_025149125\|GCF_000233455\|GCF_001078435\|GCF_000234155\|GCF_000371605\|GCF_000371585\|GCF_000371565\|GCF_000371545\|GCF_000371505\|GCF_000371485\|GCF_000424325\|GCF_000436455\|GCF_949390495\|GCF_949400645\|GCF_949474175\|GCF_949523935\|GCF_949531885\|GCF_949542115\|GCF_000371405\|GCF_001078445\|GCF_009696375\|GCF_022771995\|GCF_015549035\|GCF_005845215\|GCF_015556325\|GCF_003473545\|GCF_000954015\|GCF_013282095\|GCF_003434055\|GCF_003467385\|GCF_011317135\|GCF_013304305\|GCF_020554935\|GCF_030373645\|GCF_003466005\|GCF_902385905\|GCF_002959675\|GCF_003437595\|GCF_003458165\|GCF_003458625\|GCF_003464745\|GCF_013112035\|GCF_015547745\|GCF_015551555\|GCF_015556085\|GCF_015557875\|GCF_015558345\|GCF_016889665\|GCF_020555225\|GCF_020555685\|GCF_020559295\|GCF_020563055\|GCF_020736955\|GCF_021771435\|GCF_022137355\|GCF_022138045\|GCF_022138265\|GCF_023008325\|GCF_024460255\|GCF_024462735\|GCF_024622735\|GCF_027668515\|GCF_027671045\|GCF_030839815\|GCF_030844385\|GCF_902375545\|GCF_905197355\|GCF_958411475\|GCF_958422245\|GCF_958451465\|GCF_959023705\|GCF_959605845\|GCF_000371725\|GCF_000371705\|GCF_000371685\|GCF_000371665\|GCF_000371645\|GCF_000154365\|GCF_001078425\|GCF_949494455\|GCF_003433945\|GCF_015548445\|GCF_018785395\|GCF_020554185\|GCF_020736995\|GCF_027662505\|GCF_027666385\|GCF_900115855\|GCF_902364255\|GCF_001405335\|GCF_005844705\|GCF_006538465\|GCF_012273195\|GCF_013299965\|GCF_013304125\|GCF_013304155\|GCF_013304185\|GCF_013304205\|GCF_013304225\|GCF_013304245\|GCF_013304255\|GCF_013304275\|GCF_015152355\|GCF_015548065\|GCF_015669475\|GCF_018381395\|GCF_019012735\|GCF_020297485\|GCF_021532095\|GCF_027663445\|GCF_028210245\|GCF_028210555\|GCF_028210575\|GCF_028210655\|GCF_900100685\|GCF_900108895\|GCF_900113155\|GCF_900447015\|GCF_902374585\|GCF_910586265\|GCF_948511265\|GCF_948522125\|GCF_948524135\|GCF_948582345\|GCF_948587995\|GCF_948603615\|GCF_948681025\|GCF_948901585\|GCF_948908805\|GCF_948919595\|GCF_948932345\|GCF_948952455\|GCF_949013715\|GCF_949081035\|GCF_949121295\|GCF_958414435\|GCF_958435425\|GCF_958436665\|GCF_958453805\|GCF_959023645\|GCF_003024655\|GCF_020564305\|GCF_900102595\|GCF_902364025\|GCF_905194475\|GCF_934882645\|GCF_958421775\|GCF_018368145\|GCF_020554465\|GCF_022440645\|GCF_020709355\|GCF_020709765\|GCF_020554865\|GCF_900540675\|GCF_934402005\| |
| *Hungatella* | GCF_003201875\|GCF_027661945\|GCF_027662645\|GCF_027662785\|GCF_027696135\|GCF_932750805\|GCF_001405675\|GCF_001405995\|GCF_003435045\|GCF_003437645\|GCF_003437905\|GCF_003439535\|GCF_003466285\|GCF_003468235\|GCF_003475805\|GCF_009721605\|GCF_015553645\|GCF_015555845\|GCF_018379215\|GCF_018785385\|GCF_022834955\|GCF_022834975\|GCF_024464175\|GCF_025149285\|GCF_027671645\|GCF_902362405\|GCF_902363795\|GCF_905204275\|GCF_937936815\|GCF_959027765\|GCF_959598815\|GCF_000371445\|GCF_000433395\|GCF_000160095\|GCF_014288005\|GCF_022782185\|GCF_022784085\|GCF_014288035\|GCF_024460495\| |
| *Lachnoanaerobaculum* | GCF_003862475\|GCF_030008055\|GCF_003862485\|GCF_905371455\|GCF_001552975\|GCF_000185385\|GCF_000257705\|GCF_018372015\|GCF_000287675\|GCF_017565785\|GCF_000512995\|GCF_000296385\|GCF_003254255\|GCF_003589745\|GCF_902387945\|GCF_937890375\|GCF_938015485 |
| *Lachnoclostridium* | GCF_900078195\|GCF_000733755\|GCF_000018685\|GCF_000703105\|GCF_000702985\|GCF_905197605 |
| *Lacrimispora* | GCF_000687555\|GCF_002797975\|GCF_000526995\|GCF_000421505\|GCF_000144625\|GCF_900105615\|GCF_003432035\|GCF_000526575\|GCF_003833015\|GCF_007115105\|GCF_900155545\|GCF_009696365\|GCF_017084465\|GCF_003609635\|GCF_900205965\|GCF_000732605\|GCF_900185635\|GCF_016906045\|GCF_900105215\|GCF_900461315\|GCF_026723765 |
| ***Lachnospiraceae B*** | |
| *Agathobacter* | GCF_000020605\|GCF_002735305\|GCF_001406815 |
| *Kineothrix* | GCF_000732725\|GCF_004345255\|GCF_030863805 |
| *Acetatifactor* | GCF_003478095\|GCF_003480225\|GCF_009695995\|GCF_014337175\|GCF_024623325\|GCF_025567015\|GCF_900248245\|GCF_910584235\|GCF_910585425\|GCF_910585615\|GCF_910588225\|GCF_943193215\|GCF_947643695\|GCF_947654235\|GCF_948475165\|GCF_948475395\|GCF_948482205\|GCF_948492135\|GCF_948495795\|GCF_950096775\|GCF_950097205 |
| *Anaerobium* | GCF_900096945 |
| *Anaerocolumna* | GCF_009931695\|GCF_014218335\|GCF_018917405\|GCF_030913705\|GCF_900142215\|GCF_900205915\|GCF_947653275\|GCF_000702945\|GCF_014202875\|GCF_014218355\|GCF_029689925\|GCF_900115365\|GCF_900143645\|GCF_902479815 |
| *Anaeromicropila* | GCF_016591975 |
| *Anaerosacchariphilus* | GCF_003363435 |
| *Butyrivibrio* | GCF_025148445\|GCF_023206215\|GCF_900129945\|GCF_900143205\|GCF_900101605\|GCF_000424465\|GCF_900115735\|GCF_000145035\|GCF_000622085\|GCF_900103635\|GCF_015056685\|GCF_017433805\|GCF_017635265\|GCF_017940685\|GCF_024699075\|GCF_000424145\|GCF_900104155\|GCF_000420825\|GCF_000703165\|GCF_000424285\|GCF_000420845\|GCF_003625485\|GCF_000423945\|GCF_000421405\|GCF_000526935\|GCF_900102515\|GCF_900116875\|GCF_000621565\|GCF_000424385\|GCF_900116865\|GCF_000424585\|GCF_000621605\|GCF_000702265\|GCF_000424545\|GCF_900108105\|GCF_000420945\|GCF_000423925\|GCF_000424265\|GCF_003625475\|GCF_000420865\|GCF_900112195\|GCF_010906925\|GCF_015057245\|GCF_900218035\|GCF_000424305\|GCF_000424005\|GCF_000703005\|GCF_015057185\|GCF_017471675\|GCF_017557435\|GCF_017934625\|GCF_900100545\|GCF_900107545\|GCF_000621865\|GCF_900113655\|GCF_900104335\|GCF_008935055\|GCF_900141825\|GCF_947169445 |
| Cockroach SCG | ButyrivibrioOttesenSCG.928.D06 |
| *Eisenbergiella* | GCF_001722575\|GCF_001717125\|GCF_001717135\|GCF_001722555\|GCF_001881565\|GCF_001722585\|GCF_001722635\|GCF_001722655\|GCF_009696275\|GCF_003435265\|GCF_003435485\|GCF_003478085\|GCF_027682485\|GCF_021769355\|GCF_022781285\|GCF_027680505\|GCF_905205275\|GCF_900243045\|GCF_902385915\|GCF_902471245\|GCF_945899955\|GCF_937926915\|GCF_945830375\|GCF_945863685\|GCF_947507165 |
| *Gallintestinimicrobium* | GCF_025567525.1 |
| *Roseburia* | GCF_025567465\|GCF_000225345\|GCF_900537995\|GCF_000174195\|GCF_014287435\|GCF_009695765\|GCF_014287515\|GCF_003612565\|GCF_001940165\|GCF_014297335\|GCF_014287635.1 |
| *Suilimivivens* | GCF_003612395\|GCF_009917485\|GCF_025567405.1 |
| ***Lachnospiraceae C*** | |
| *Lachnospira* | GCF_000146185\|GCF_000424105\|GCF_000702205\|GCF_003458705\|GCF_009680455\|GCF_014287955\|GCF_020564355\|GCF_900103815 |
| *Mediterraneibacter* | GCF_000153925\|GCF_001487105\|GCF_014287475\|GCF_016902345\|GCF_020687545\|GCF_025152405\|GCF_900120155\|GCF_934309345\|GCF_934309415\|GCF_934330165\|GCF_944377805\|GCF_003574295 |
| *Anaerosporobacter* | GCF_012070565\|GCF_900142955 |
| *Anaerostipes* | GCF_002270485\|GCF_005280655\|GCF_014467075\|GCF_016586355\|GCF_018381315\|GCF_018918155\|GCF_018982945\|GCF_025567365\|GCF_030296915 |
| *Blautia* | GCF_001689125\|GCF_002222595\|GCF_002270465\|GCF_003287895\|GCF_013304445\|GCF_014287615\|GCF_900120295\|GCF_900461125\|GCF_947654165 |
| *Coprococcus* | GCF_003482105\|GCF_025149915\|GCF_025567285\|GCF_025567345\|GCF_019734885\|GCF_902381825 |
| *Cuneatibacter* | GCF_004216775 |
| *Dorea* | GCF_025150245 |
| *Extibacter* | GCF_001185345\|GCF_004345005\|GCF_008281175 |
| *Hominisplanchenecus* | GCF_020687205\|GCF_943193015 |
| Cockroach SCG | Lachnospiraceae bacterium OttesenSCG-928-E19\|Lachnospiraceae bacterium OttesenSCG-928-J05 |
| *Lachnotalea* | GCF_003201285\|GCF_008830185\|GCF_900184995 |
| *Metalachnospira* | GCF_018918145 |
| *Pseudolachnospira* | GCF_022867805 |
| *Robinsoniella* | GCF_000797495 |
| *Sellimonas* | GCF_001280875\|GCF_019754295\|GCF_027924685 |

**Supplemental Table 4: Glycoside hydrolases associated with cellulose or xylan degradation.** Adapted from dbCAN2 substrate chart.

| **Substrate** | **Family** | **Description** | **Additional substrates** |
| --- | --- | --- | --- |
| **Xylan** | **GH1** | β-xylosidase | β-fucosides, β-galactan, β-glucan, β-glucuronan, β-mannan |
|  | **GH10** | endo-1,3-β-xylanase; endo-1,4-β-xylanase; arabinoxylan-specific endo-β-1,4-xylanase | β-glucan |
|  | **GH11** | exo-1,4-β-xylosidase; endo-β-1,4-xylanase |  |
|  | **GH115** | xylan ⍺-1,2-glucuronidase |  |
|  | **GH120** | β-xylosidase |  |
|  | **GH141** | xylanase | pectin |
|  | **GH2** | β-xylosidase | ⍺-mannan, arabinan, β-galactan, β-glucan, β-glucuronan, β-mannan, chitosan, host glycan, pectin |
|  | **GH26** | β-1,3-xylanase | β-glucan, β-mannan |
|  | **GH3** | xylan 1,4-β-xylosidase | arabinan, β-glucan, chitin, host glycan, peptidoglycan |
|  | **GH30** | β-xylosidase; endo-β-1,4-xylanase; glucuronoarabinoxylan endo-β-1,4-xylanase | β-fucosides, β-glucan, β-glucuronan, host glycan |
|  | **GH4** | ⍺-glucuronidase | β-glucan, pectin, starch, sucrose |
|  | **GH43** | β-1,3-xylosidase; β-xylosidase; ⍺-L-arabinofuranosidase; xylanase | arabinan, β-galactan |
|  | **GH51** | endo-β-1,4-xylanase | arabinan, β-glucan |
|  | **GH67** | xylan ⍺-1,2-glucuronidase; ⍺-glucuronidase |  |
|  | **GH95** | ⍺-L-galactosidase | xyloglucan, host glycan, pectin |
|  | **GH98** | endo-β-1,4-xylanase | host glycan |
| **Cellulose** | **GH48** | reducing end-acting cellobiohydrolase; endo-β-1,4-glucanase | chitin |
|  | **GH74** | endoglucanase | xyloglucan |
|  | **GH9** | endoglucanase; exo-β-1,4-glucanase / cellodextrinase; cellobiohydrolase | β-glucan, chitosan |
|  | **GH94** | cellobiose/cellodextrin/cellobionic acid phosphorylase | β-glucan, chitin |
| **Xylan & Cellulose** | **GH39** | exo-β-1,4-glucanase / cellodextrinase | β-galactan, β-glucan, host glycan |
|  |  | β-xylosidase; ⍺-L-arabinofuranosidase |  |
|  | **GH5** | exo-β-1,4-glucanase / cellodextrinase; cellulose β-1,4-cellobiosidase | arabinan, β-glucan, β-glycan, β-mannan, chitin, chitosan |
|  |  | arabinoxylan-specific endo-β-1,4-xylanase |  |
|  | **GH8** | cellulase | β-glucan, chitosan |
|  |  | reducing-end-xylose releasing exo-oligoxylanase; endo-1,4-β-xylanase |  |

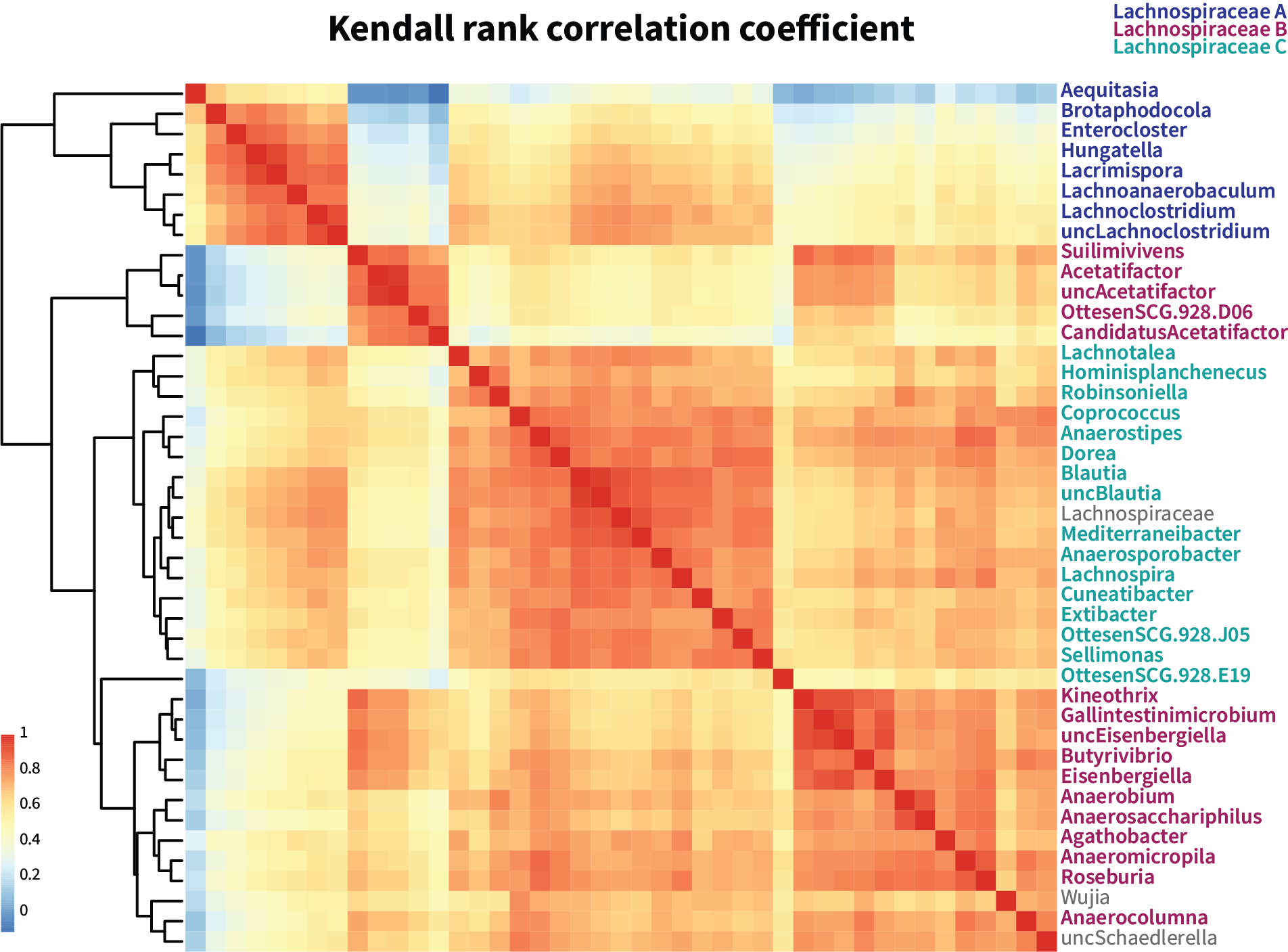

**Supplemental Figure S1:** Kendall Tau was calculated to identify co-occurrence of abundant genera within Lachnospiraceae. Co-occurring genera were grouped into pangenomes A, B, or C.

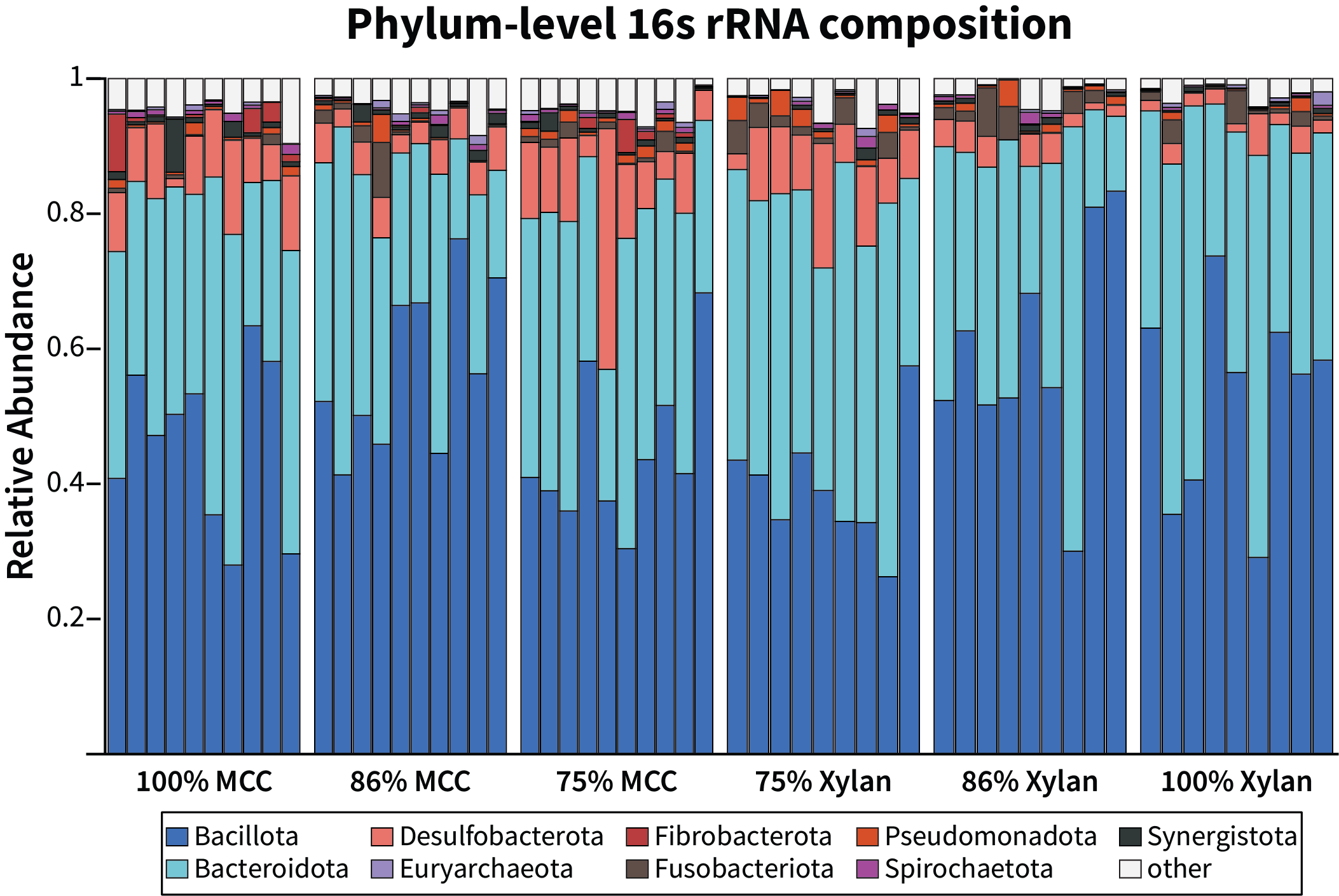

**Supplemental Figure S2: Phylum-level community composition of cockroaches fed xylan differ from those fed cellulose.** ASV count tables were aggregated at the phylum taxonomic level and converted to proportions. Phyla that comprised at least 1% of one sample were kept for visualization, while low abundant phyla were collapsed into “Other”. MCC = microcrystalline cellulose

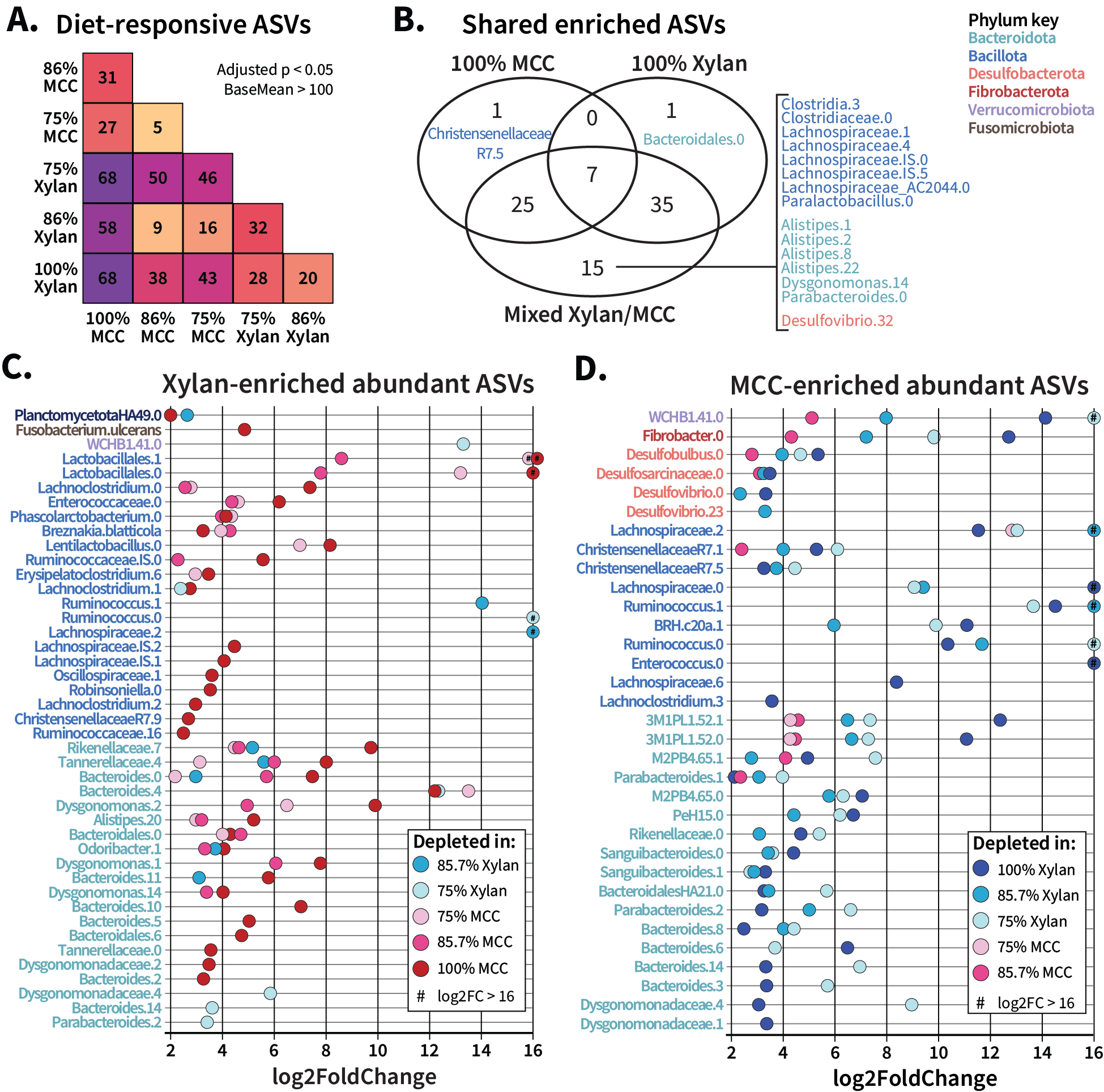

**Supplemental Figure S3: Diet-based enrichment of ASVs on mixed and pure fiber diets.** Enrichment of ASVs on the ratio diets was assessed with DESeq2, and pairwise results were extracted. (**A**) The total number of abundant (baseMean > 100) significantly differentially abundant ASVs identified in pairwise comparisons were plotted as a heatmap matrix, with color scaled within the total heatmap. (**B**) ASVs found to be enriched in 100% xylan, 100% cellulose, and any of the ratio diets were compared as a Venn diagram to identify overlap; since the ratio diets were aggregated, some ASVs appeared as significant in all three circles. (**C**) All abundant ASVs enriched by xylan were plotted against their log2 fold change vs the other five diets (pairwise comparison indicated by color). (**D**) All abundant ASVs enriched by cellulose were plotted against their log2 fold change vs the other five diets. MCC = microcrystalline cellulose; # =log2FC > 16

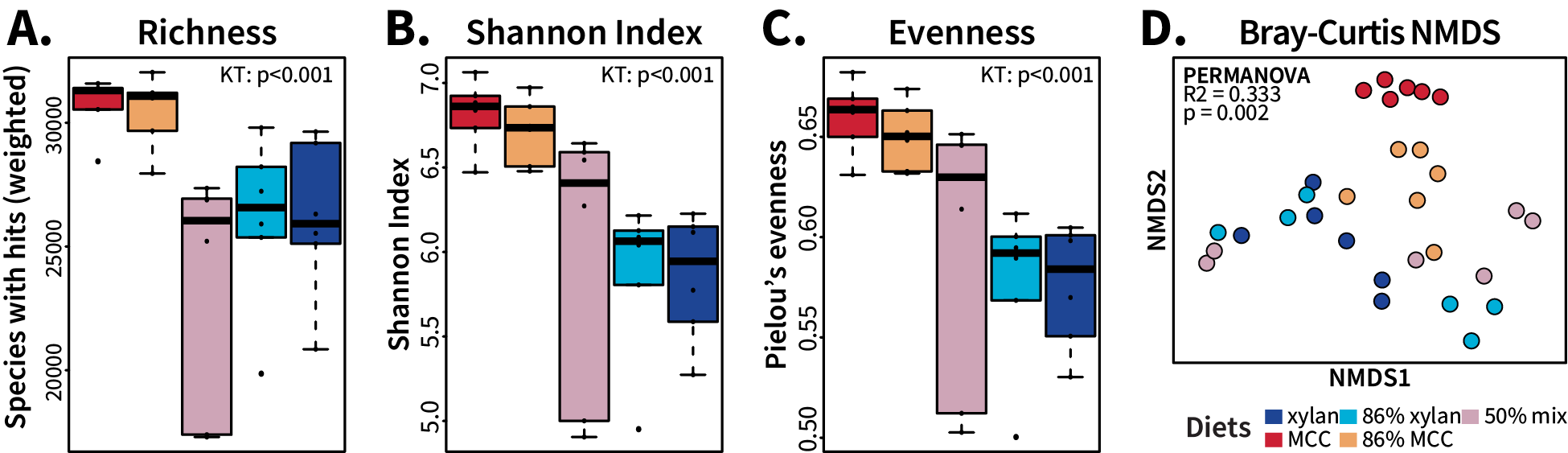

**Supplemental Figure S4: Alpha and beta diversity of taxonomic composition of metatranscriptomes.** The total counts of taxonomic identity assigned to the transcripts within each metatranscriptome was rarefied to 4,740,617 prior to comparing the alpha diversity measures (A) richness, (B) Shannon index, and (C) evenness as well as (D) Bray-Curtis dissimilarity of the different fiber diets. Significance of the alpha diversity tests was determined with Kruskal-Wallis test. Bray-Curtis dissimilarity ordination was produced by NMDS and evaluated for significance with PERMANOVA.

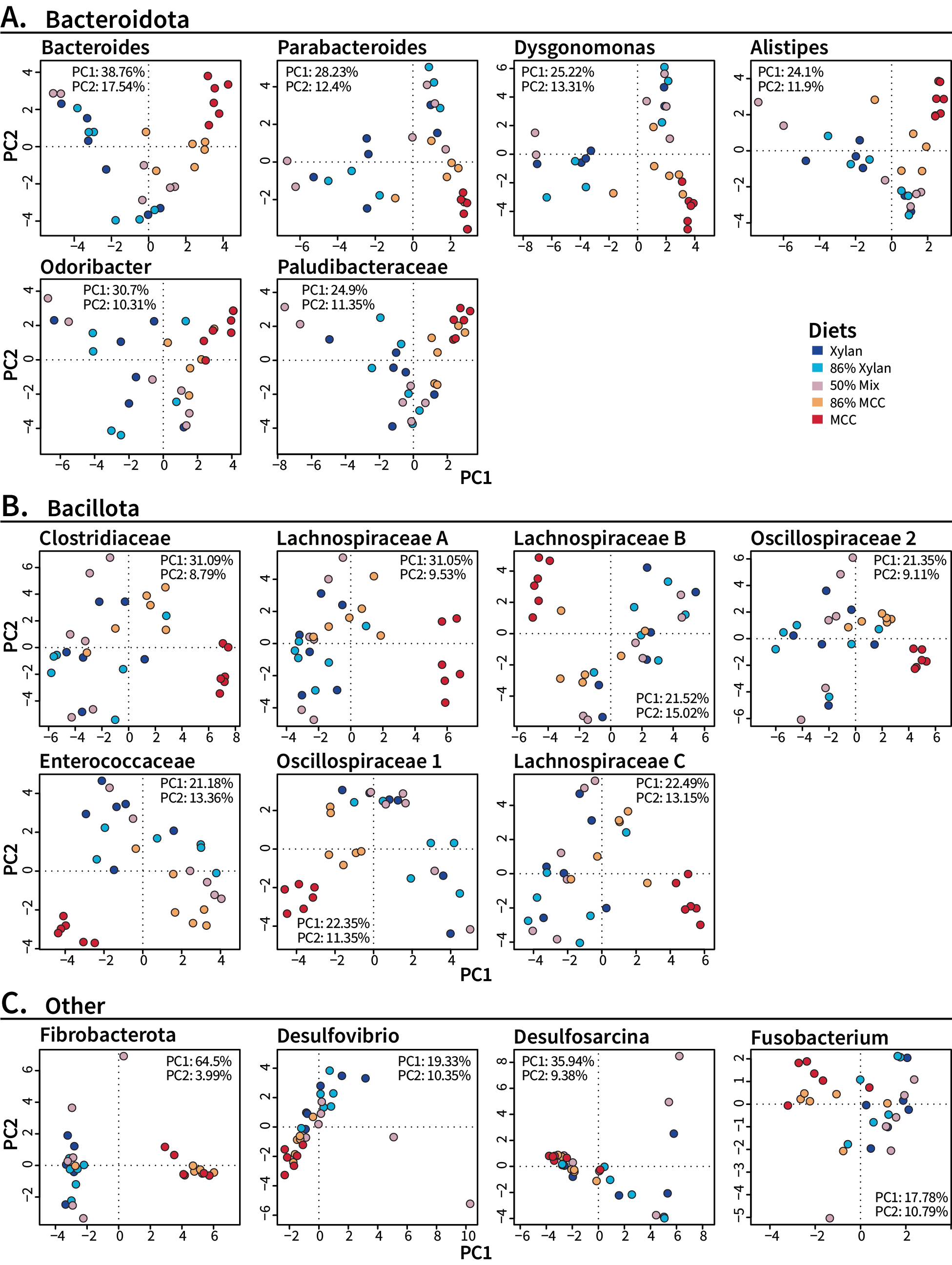

**Supplemental Figure S5: Principal Component Analysis of gene cluster expression within each pangenome.** Gene cluster expression for pangenomes from (**A**) *Bacteroidota,* (**B**) *Bacillota,* and (**C**) other phyla was normalized using the variance-stabilizing transformation provided with DESeq2 prior to unconstrained principal component calculations and subsequent ordination.

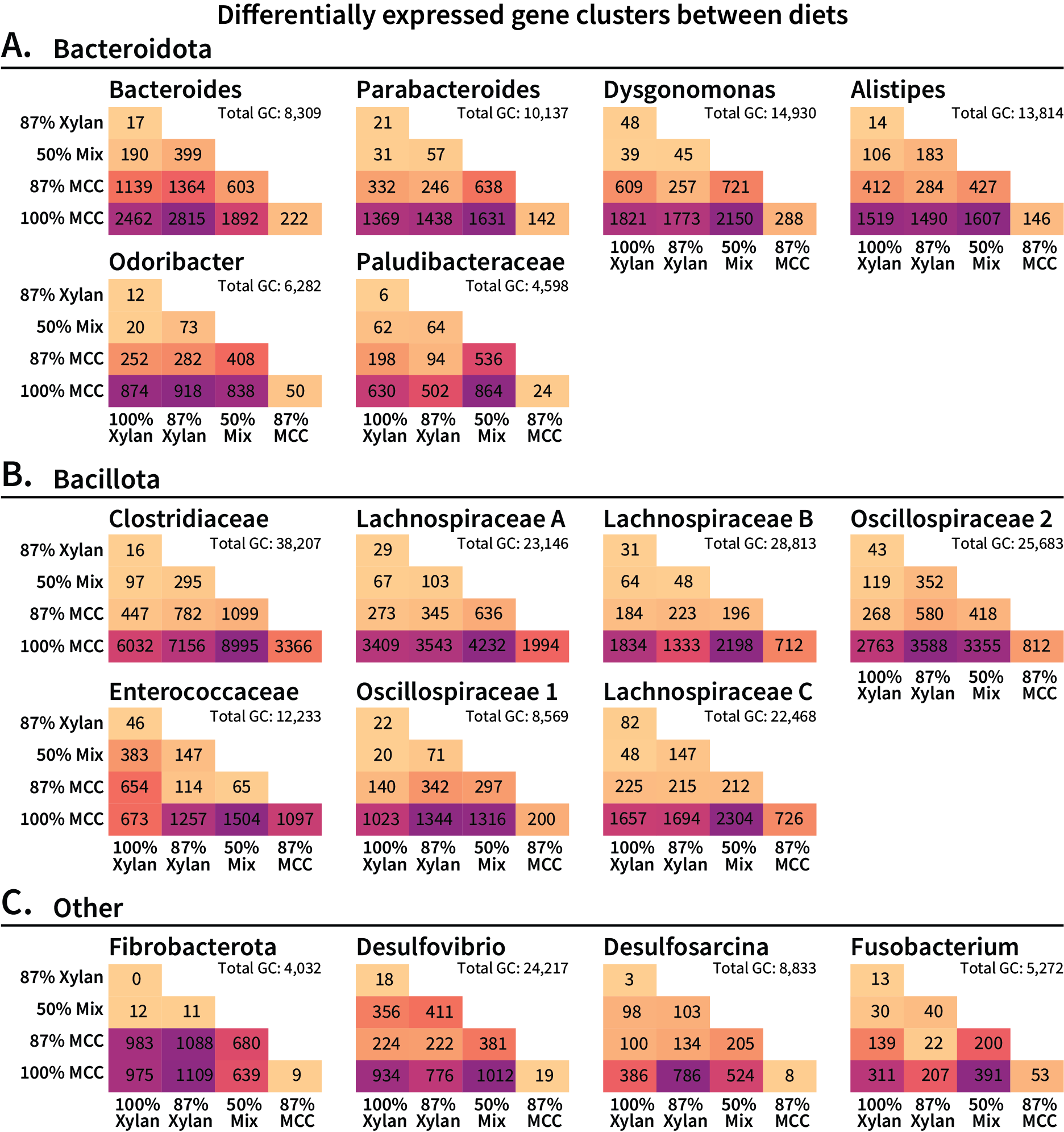

**Supplemental Figure S6: Pangenome-specific differentially expressed gene clusters between fiber diets.** Gene cluster count matrices were analyzed with DESeq2, fit to local sample dispersion. Pairwise results were pulled out using “contrast”, and the total number of differentially expressed gene clusters with padj > 0.05 were summed for plotting. Heatmap color is scaled within each pangenome. GC: gene cluster.

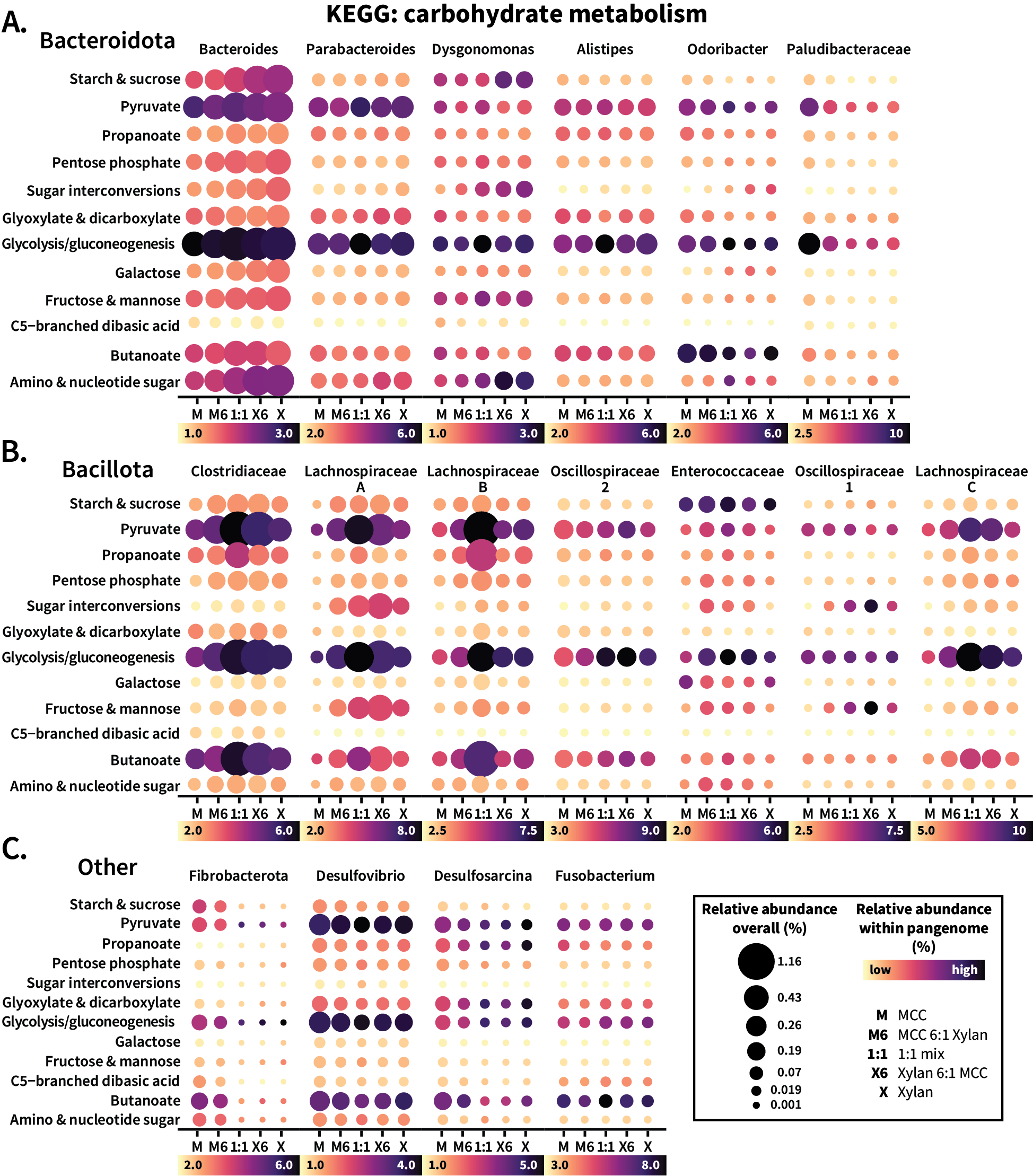

**Supplemental Figure S7: Patterns of carbohydrate-degrading metabolic pathways across a gradient of fiber ratios.** Pangenomes from (**A**) *Bacteroidota*, (**B**) *Bacillota*, and (**C**) other phyla were analyzed for KEGG carbohydrate metabolic pathway enrichment across the pure- and mixed-fiber diets. Gene clusters annotated with KEGG orthologs belonging to relevant metabolic pathway were summed together, divided by both pangenome transcripts (color) and total transcripts (size) per sample, then averaged by diet for plotting. Size is scaled based on total transcriptional abundance and therefore uses the same key for each pangenome, while color is scaled within each pangenome according to the key beneath each plot. M: 100% microcrystalline cellulose; M6: 86% cellulose; 1:1: 50% xylan and 50% cellulose; X6: 86% xylan; X: 100% xylan.

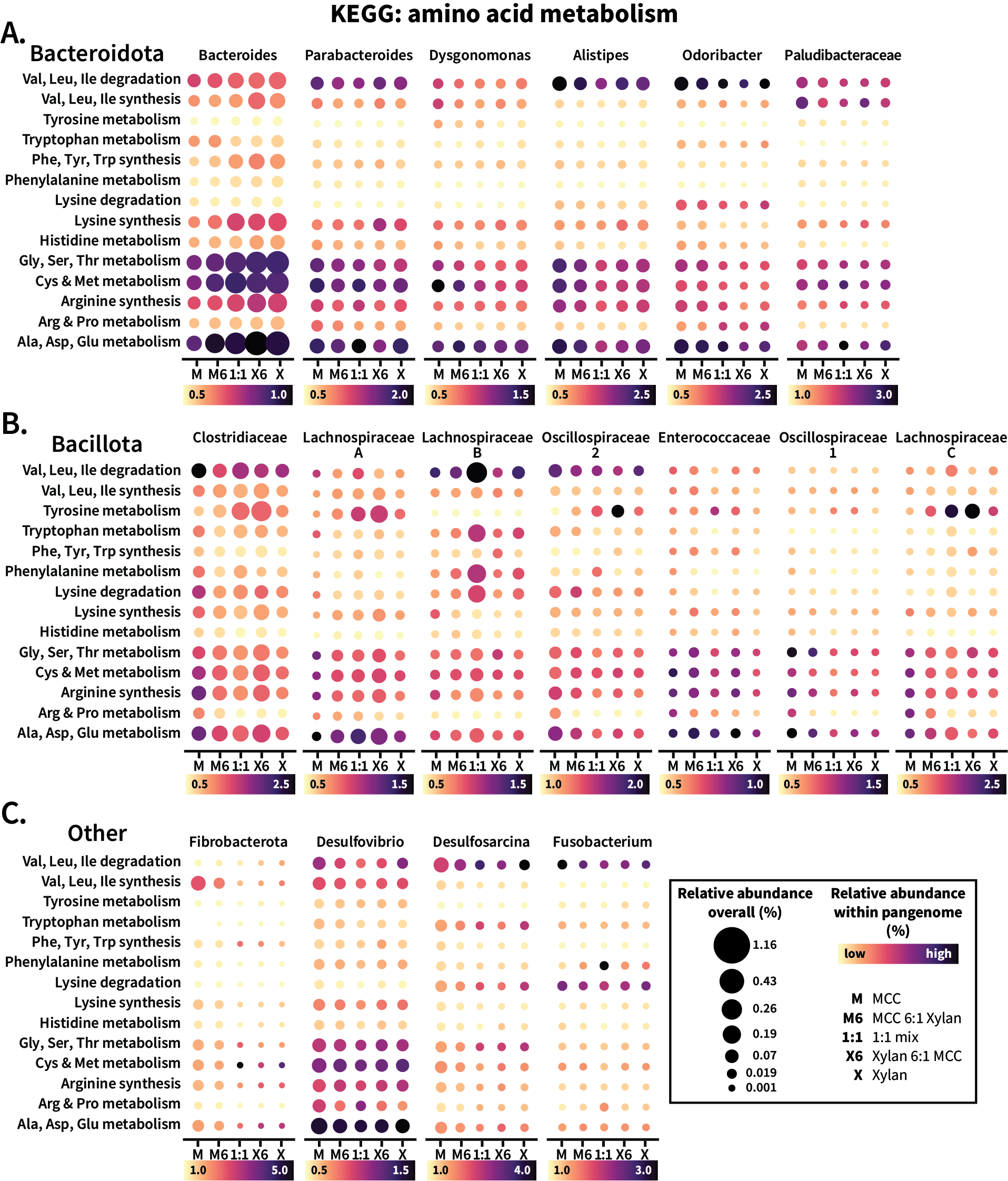

**Supplemental Figure S8: Patterns of amino acid processing metabolic pathways across a gradient of fiber ratios.** Pangenomes from (**A**) *Bacteroidota*, (**B**) *Bacillota*, and (**C**) other phyla were analyzed for KEGG amino acid metabolic pathway enrichment across the pure- and mixed-fiber diets. Gene clusters annotated with KEGG orthologs belonging to each relevant metabolic pathway were summed together, divided by both pangenome transcripts (color) and total transcripts (size) per sample, then averaged by diet for plotting. Size is scaled based on total transcriptional abundance and therefore uses the same key for each pangenome, while color is scaled within each pangenome according to the key beneath each plot. M: 100% microcrystalline cellulose; M6: 86% cellulose; 1:1: 50% xylan and 50% cellulose; X6: 86% xylan; X: 100% xylan.

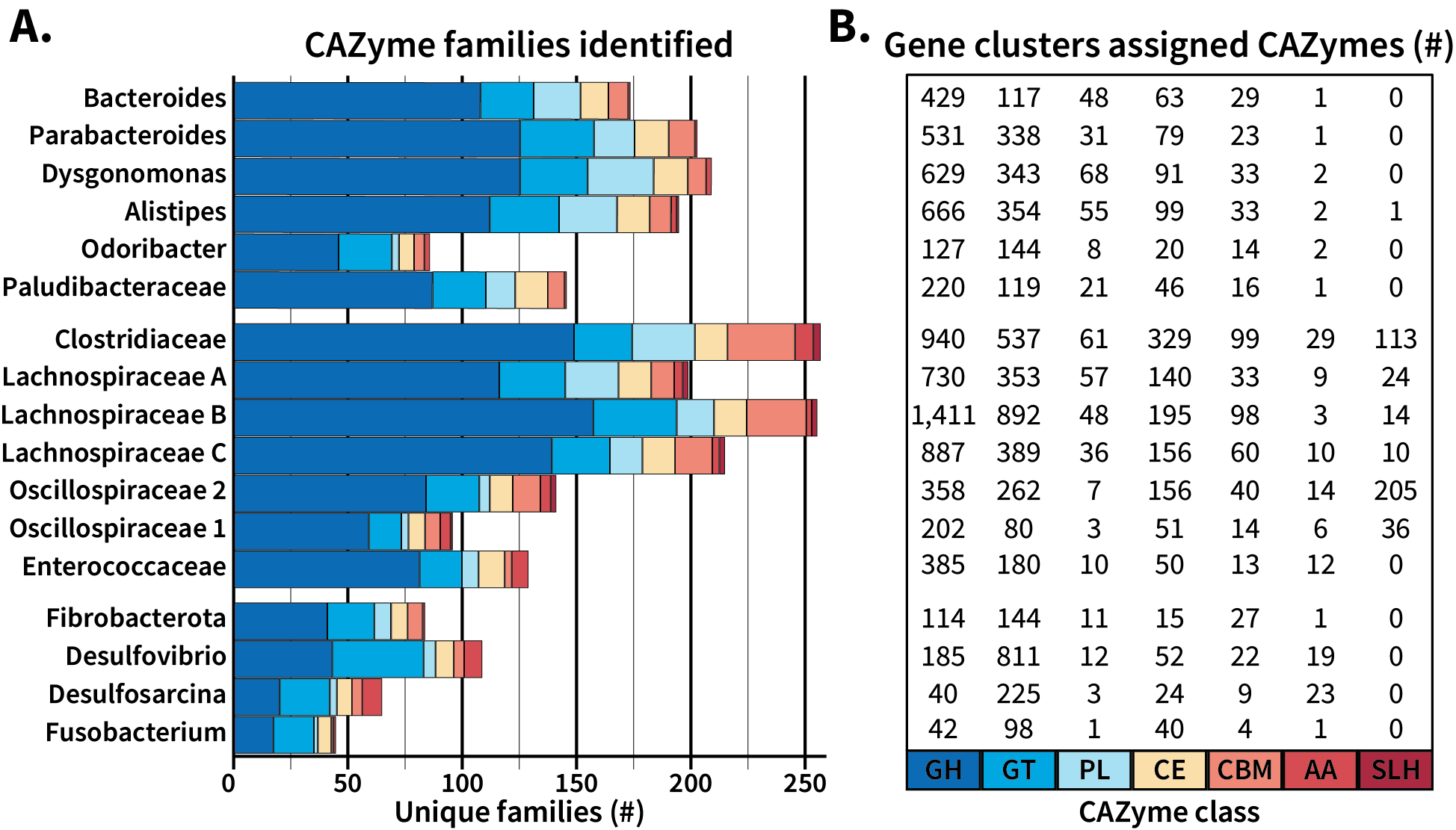

**Supplemental Figure S9: CAZyme family type and distribution expressed by the pangenomes. (A)** The total number of unique CAZyme families found in the pangenomes assessed is plotted, colored by CAZyme class. **(B)** The total number of gene clusters annotated as each class. GH: glycoside hydrolase; GT: glycosyl transferase; PL: polysaccharide lyase; CE: carbohydrate esterase; AA: auxiliary activity; CBM: carbohydrate binding module; SLH: S-layer homology domain

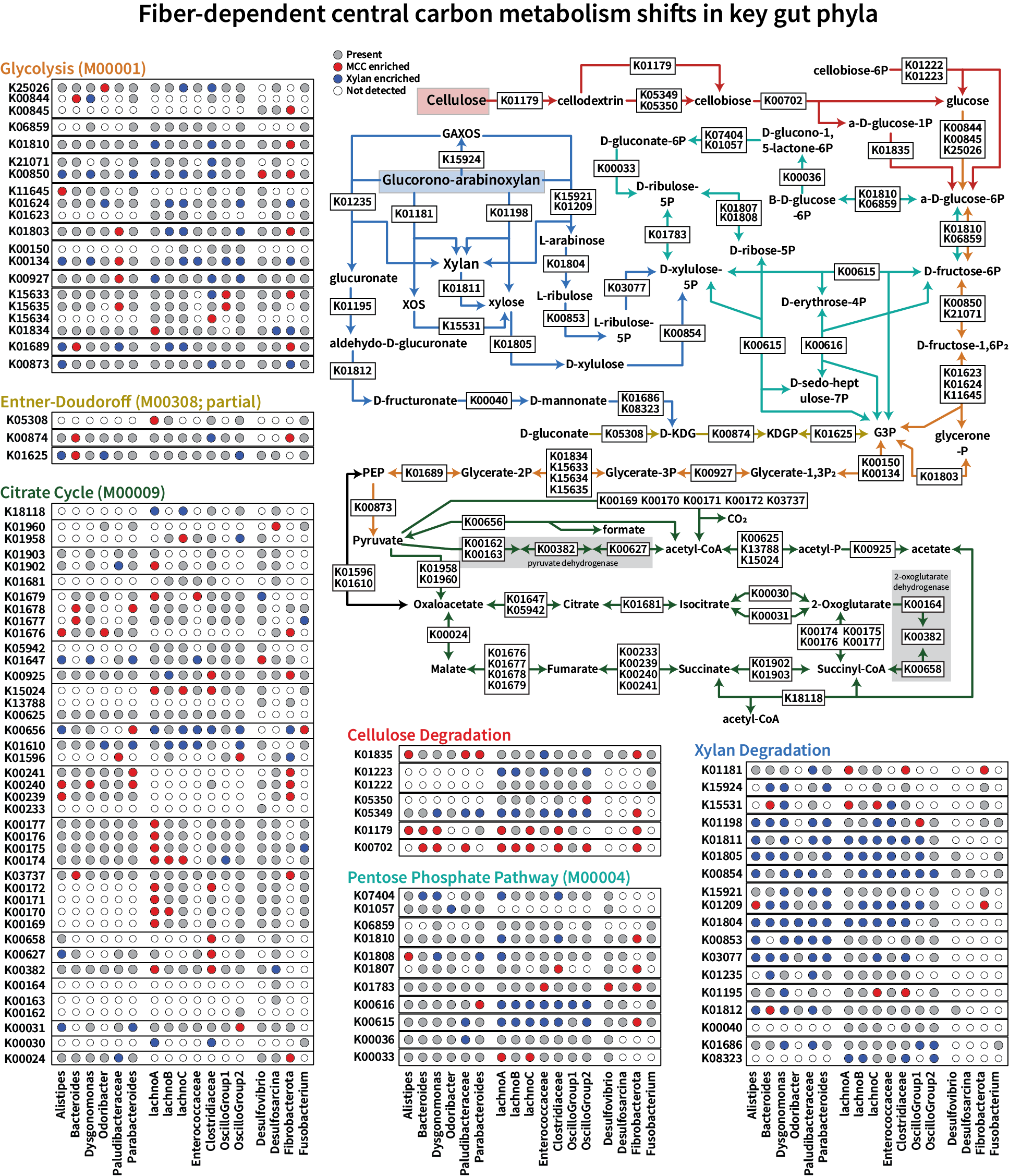

**Supplemental Figure S10: KEGG map and signficance by step of central carbon metabolism in cellulose and xylan degradation.**

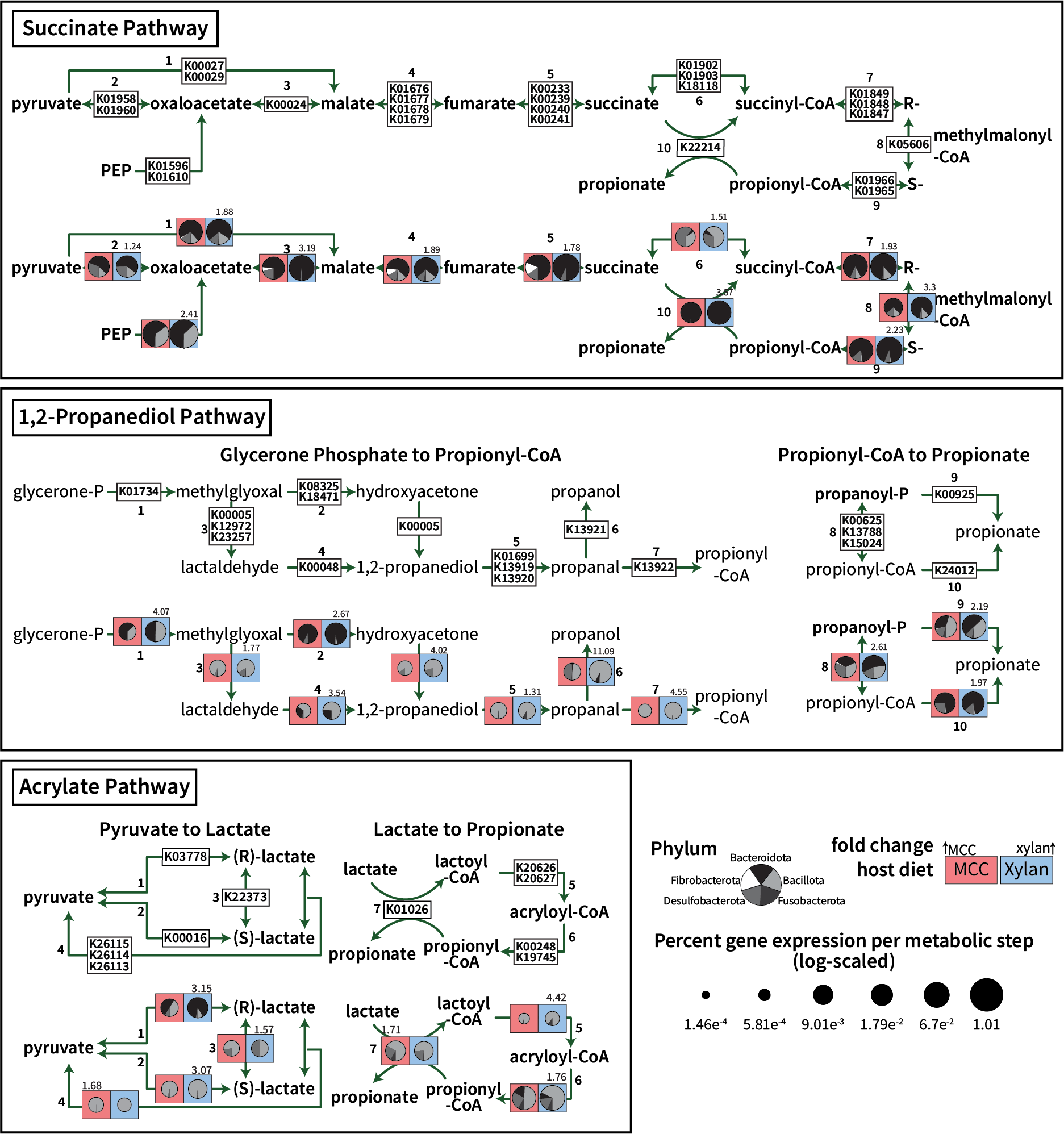

**Supplemental Figure S11: Propionate production pathways in response to fiber.** Three pathways of propionate production are presented, with pie charts created as described in Figure S5. The size of the pie charts indicates the overall transcriptional percent that step comprises in the 100% cellulose (red box) and 100% xylan (blue box) diets, with the number indicating the fold increase in relative abundance in the direction of either cellulose (left corner) or xylan (right corner). Pathways are indicated with arrow color, and the transparent boxes are included to focus attention on xylan degradation steps (blue) and cellulose degradation steps (red).
